# Biophysics-informed generative models improve activity and thermostability of designed ribozymes

**DOI:** 10.64898/2026.11.21.768297

**Authors:** Francesco Calvanese, Lorenzo Rosset, Polina Pavlinova, Louis Meuret, Vaitea Opuu, Samuela Pasquali, Philippe Nghe, Francesco Zamponi, Jorge Fernandez-de-Cossio-Diaz, Martin Weigt

## Abstract

Generative models of biological sequences can produce molecules that fold and function similarly to natural ones. A further challenge is to steer generation toward a desired property, such as resistance to high temperature. This is particularly difficult when available sequence data are limited, and typically requires probing thousands of variants experimentally, either through directed evolution or to retrain a model specialized to the targeted property. Ideally, however, one would avoid such an expensive experimental step. Here, we show that biophysical information can complement limited evolutionary data to guide the design of catalytic RNA molecules toward increased activity and thermostability. We combine a sequence generative model with a physical score based on the Turner nearest-neighbor model, whose parameters were originally derived from calorimetric measurements, independently of the sequence family considered. On its own, this physical score is ineffective for design: optimizing it destroys function, likely because it is blind to the catalytic core and to tertiary contacts. However, when coupled to the generative model, it provides complementary information that enables better designs than evolutionary data alone. We demonstrate this on self-splicing group I intron ribozymes: starting from designs that are active at 37 °C but inactive at 60 °C, fewer than ten substitutions are sufficient to confer catalytic activity at 60 °C.

## I. INTRODUCTION

Generative models of biological sequences can produce molecules that adopt the structures and functions characteristic of natural families [1–4]. A key next step is to harness these models for controlled engineering: enhancing a desired property while preserving the constraints essential for biological function. Because unconditioned generation tends to reproduce the range of properties found in natural sequences, targeted engineering requires steering the learned distribution toward a desired property without leaving the space of evolutionarily plausible, functionally viable molecules. This challenge becomes particularly acute when evolutionary sequence data are limited. In this regime, increasingly large and expressive family-specific models cannot simply be trained on more data, motivating the use of complementary sources of information to improve sequence design.

We address a concrete instance of this problem in self-splicing group I intron ribozymes. These RNAs catalyze their own excision from a precursor transcript through two sequential transesterification reactions, which require a conserved catalytic core held together by long-range tertiary contacts [5, 6]. Our starting sequences, generated previously by a standard generative model based on evolutionary sequence data alone, splice efficiently at 37 °C but show no detectable activity at 60 °C [2].

Extending biomolecular catalytic activity to elevated temperatures is an important engineering objective in many practical applications, as higher operating temperatures can accelerate reactions, reduce contamination, and facilitate integration with other processes. At the same time, we seek to remain as close as possible to the starting sequence so as to preserve its existing functional properties and avoid unintended changes to an already optimized molecular background. We therefore impose a limited mutational budget.

Existing approaches to property-guided sequence engineering broadly include directed evolution, optimization based on supervised predictors, and alignment of generative models with predictive scores [7]. Directed evolution identifies improved variants by synthesizing and screening large libraries, a process that is effective but must generally be repeated for each molecular system and target property. Supervised predictors learn mutational effects from characterized variants and are far cheaper to query. However, optimizing sequences directly against their predictions can drive the design outside the training distribution, potentially disrupting conserved motifs, structural interactions, or other epistatic constraints that are necessary for function. Model-alignment methods mitigate this risk by combining the predictive signal with a generative prior over plausible sequences [8–10]. What these approaches share is a dependence on a sufficiently large, target-specific set of experimentally characterized variants, whether they are screened directly or used to fit a predictor. This dependence is particularly restrictive for RNA, for which sequence-to-function datasets remain considerably scarcer and smaller than for proteins. Moreover, evolutionary sequence families themselves can be relatively small, limiting how much family-specific information can be extracted by highly parameterized generative models. In such data-limited settings, improving the model need not mean increasing its complexity; it can instead mean supplying additional information from an independent source.

We circumvent the need for target-specific labeled data by deriving the design signal from a physical model. RNA secondary-structure thermodynamics provides a natural source of such complementary information. In particular, the Turner nearest-neighbor model distills decades of calorimetric measurements into a semi-empirical description of RNA folding energetics, allowing sequence-dependent properties related to structure formation and thermal stability to be evaluated at low computational cost [11]. Because its parameters describe local stacking and loop energetics arising from nucleotide chemistry, rather than features learned from a particular sequence family, the model can be applied across families without retraining and is thus not impacted when family-specific data are limited. However, it does not capture the family-specific constraints required for biological function. Consequently, using the thermodynamic signal alone could favor sequences with improved folding energetics that no longer retain catalytic activity.

We therefore combine this physical signal with family-specific information inferred from evolution. A direct coupling analysis (DCA) model [12] captures the sequence constraints characteristic of the natural family and defines a statistical prior over family-compatible sequences. Its relatively compact parameterization makes it particularly suitable for learning from evolutionary datasets of limited size [13, 14]. We augment this prior with a term derived from the Turner model that favors greater thermal robustness, integrating the two sources of information into a single statistical score for sequence generation [15]. Intuitively, the two terms provide complementary phenotypic information: the evolutionary model captures family-specific constraints needed for function, whereas the physical model contributes trans-ferable information about RNA folding and thermal stability. Sampling from the resulting distribution therefore steers generation toward sequences with improved thermal properties while maintaining the evolutionary constraints associated with functional family members. In this way, biophysical knowledge effectively supplements information derived from evolutionary data alone.

High-throughput screening of 2,500 variants derived from 10 starting sequences shows that most designs generated by the combined model recover robust self-splicing activity at 60 °C. Neither the evolutionary model nor the thermodynamic score alone yield comparable activity, demonstrating that successful design depends on coupling the two sources of information. Notably, activity at 60 °C is achieved with only 5–10 substitutions, corresponding to 2.5%–5% of the sequence. These results show that embedding a biophysical score within a family-specific generative prior transforms it into an effective signal for sequence engineering. More broadly, this approach enables property-directed generation without a labeled dataset for the target family or iterative experimental feedback during the design process.

## II. RESULTS AND DISCUSSION

### A. An effective Hamiltonian for MNIST

We introduce the framework using a toy system in which plausibility and the target property can be inspected visually. We consider the subsets of zeros and eights in the binarized MNIST dataset [16]. Each configuration is a 28 × 28 binary image, *x ∈* {0, 1} ^784^, with label *y ∈* {0, 8}. Starting from an image of a zero, *x*^st^, the task is to transform it into a plausible eight, *x*^mut^, using a limited number of spin flips, quantified by the Hamming distance *d*_*H*_ (*x*^st^, *x*^mut^). The digit label represents the target property, which we score using a logistic regression classifier *p*(*y* |*x*). To follow a physics-inspired energy convention in which favorable configurations have low values, we define the steering potential as

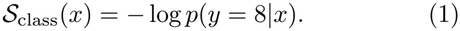

Thus, *S*_class_(*x*) is low for configurations that the classifier identifies as eights and high for configurations identified as zeros.

For the 0-to-8 task, the classifier relies strongly on the central pixels, which tend to be active in eights and in-active in zeros. Minimizing *S*_class_ under a limited pixel budget therefore fills the center of the zero, producing images that are confidently classified as eights but remain visually incoherent. The classifier favors the target label without enforcing the structure of a valid digit.

We model the distribution of valid digits with a Restricted Boltzmann Machine (RBM). The RBM is trained in an unsupervised manner on the combined set of zeros and eights. Its role is to learn the statistical structure of the images in the training distribution. The RBM defines a probability distribution over the space of pixel configurations that can be written in Boltzmann form,

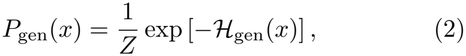

where *ℋ*_gen_(*x*) is the statistical energy, or Hamiltonian, assigned to *x*, and *Z* is the normalizing partition function. Configurations consistent with the training data occupy low-energy regions of *ℋ*_gen_, whereas irregular pixel arrangements are assigned higher energies.

Our goal is to devise a sampling strategy that controls generative plausibility, quantified via *ℋ S*_gen_(*x*), while steering toward the target property, measured by _class_(*x*), and limiting the mutational distance *d*_*H*_ (*x*^st^, *x*) from the starting sequence. According to the maximum-entropy principle, the distribution with prescribed expected values of these quantities is governed by the effective Hamiltonian

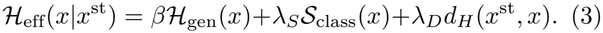

The Lagrange multipliers *β, λ*_*S*_, and *λ*_*D*_ set the relative strengths of the three constraints. With positive *λ*_*S*_, minimizing the effective Hamiltonian favors low values of *S*_class_ and therefore configurations with a high probability of belonging to the target class [17]. A Bayesian interpretation of this procedure as tempered conditional sampling is given in the Supplementary Information (SI, Sec. S1).

To illustrate the roles of the generative and steering terms, we fix a starting zero from MNIST and a target distance *D*, scan *β* and *λ*_*S*_, and tune *λ*_*D*_ to keep most samples near *D*. Fig. 1 maps the resulting configurations in the plane (ℋ_gen_, *S*_class_). Samples driven primarily by the classifier (magenta dots) occupy the noisy-eight regime, characterized by low steering potential and high generative energy. Increasing *β* suppresses these configurations and shifts sampling toward recognizable digits. Plausible eights occupy the region in which both quantities are low (orange dots), and plausible zeros are found at low generative score and high steering score (blue dots). This observation shows visually how the generative prior constrains optimization of the property score.

**FIG 1.**
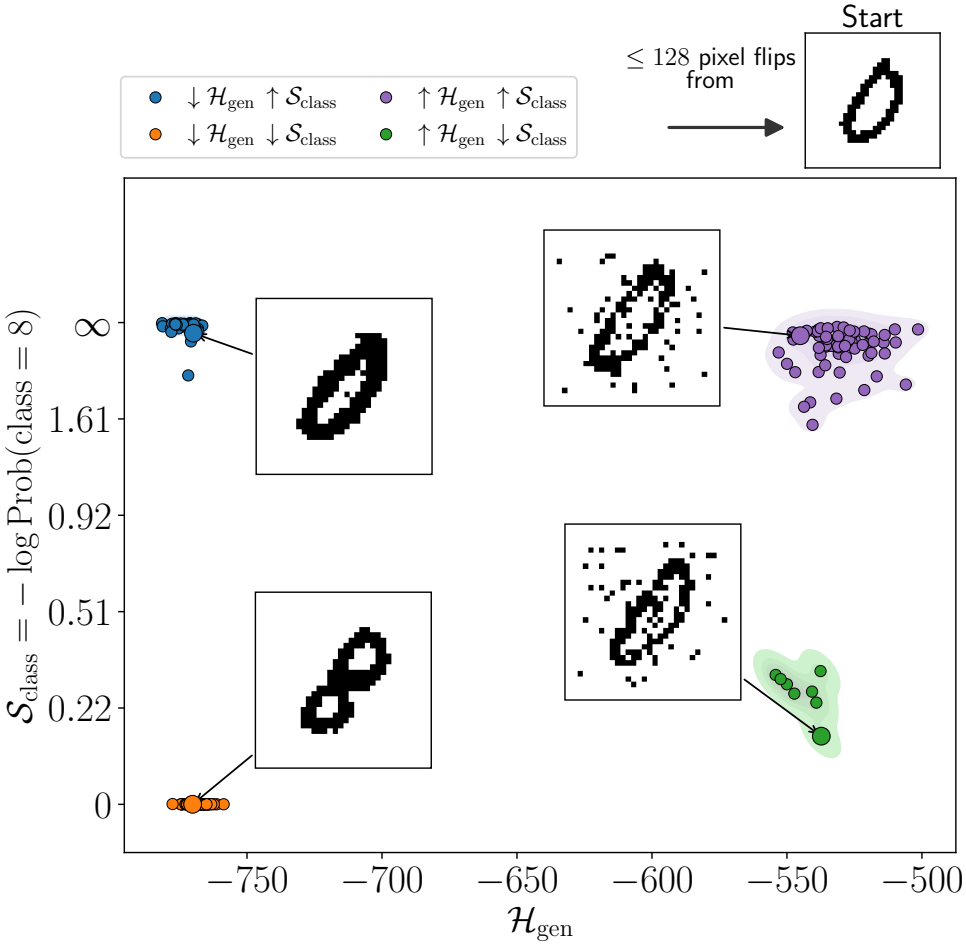
Property-directed sampling in the MNIST 0-to-8 transformation. Samples generated at a limited Hamming distance budget (≤ 128 pixel flips) from the starting zero are shown in the plane defined by the generative energy ℋ_gen_(x) and the steering potential S_class_(x). Low steering potential corresponds to a high probability of the target class. The four regimes correspond to valid zeros (↓ ℋ_gen_, ↓ S_class_), valid eights (↓ ℋ_gen_, ↓ S_class_), noisy zeros (↑ ℋ_gen_, ↑ S_class_), and noisy eights (↑ ℋ_gen_, ↓ S_class_).

### B. Designing introns for high-temperature self-splicing

Eq. (3) provides the template for ribozyme design. In the following sections, the generative and steering terms are specified using a DCA model of the group I intron family and a thermodynamic score derived from the Turner model, respectively.

The effective-Hamiltonian formulation in Eq. (3) is particularly natural for energy-based generative models, which assign an energy to complete configurations and sample them using Markov Chain Monte Carlo methods [18]. At each Monte Carlo step, the generative energy, property score, and mutational distance can be computed directly from the complete configuration and combined in the sampling energy. Autoregressive and diffusion models operate on incomplete or noised representations of the data, for which global quantities such as mutational distances and target-property proxies are not directly available [19]. Incorporating these quantities therefore requires additional procedures such as guided decoding, reconstruction-based estimates, or post-generation rescoring [20] [21, 22].

Group I introns are catalytic RNAs that catalyze their own excision from a precursor RNA and join the flanking exons. This process, known as self-splicing, requires the RNA to adopt a precise three-dimensional structure stabilized by conserved interactions between distant parts of the sequence. These ribozymes are well suited to sequence-design studies because their activity can be measured across large variant libraries using high-throughput assays, which exploit the fact that they catalyze their own splicing and have little crossreactivity. They are also of broader scientific interest because some group I introns can assemble into active ribozymes from inactive RNA fragments and form cooperative catalytic networks. These properties have made them useful models for studying RNA self-organization and molecular cooperation in origin-of-life research [23, 24].

Our starting data come from Lambert et al. [2], who used several generative models, including DCA, variational autoencoders (VAE) and covariance models, to produce sequence variants based on the *Azoarcus* group I intron. The generated sequences retained the full construct length of 197 nucleotides, of which 193 were modeled by DCA while spanning a broad range of mutational distances from wildtype. Self-splicing activity was measured for tens of thousands of designed variants at 37 °C and 60 °C. Activity at 60 °C was strongly conditioned on activity at 37 °C (only 20 of the 2292 variants active at 60 °C were inactive at 37 °C) and decreased with mutational distance from the wild type (Fig. S1). Refer to Sec. S6.6 and Fig. S6 for the definitions of activity cut-offs.

From this dataset, we selected ten variants carrying between 25 and 30 substitutions with respect to the *Azoarcus* wildtype. These variants are chosen to splice efficiently at 37 °C, but they show no detectable activity at 60 °C (see Sec. S2 and Table S1). For each starting variant, our goal is to design sequences containing 5 to 10 additional substitutions that recover activity at 60 °C while retaining activity at 37 °C.

The identities, selection criteria, and baseline properties of the ten starting sequences are detailed in the SI (Secs. S2.1 and S2.2).

Applying the effective-Hamiltonian framework requires a family-specific term that measures sequence plausibility and a steering term associated with thermal robustness. We define these components in the following sections.

### C. An evolutionary prior for functional sequences

We quantify sequence plausibility using the DCA model of Lambert et al. [2]. The model was trained on a multiple sequence alignment (MSA) of *M* = 817 homologous group I introns of aligned length *L*, denoted by 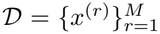, where 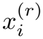 is one of the four nucleotides or an alignment gap. The alignment places homologous (i.e. evolutionarily corresponding) residues in the same columns, allowing the model to learn both nucleotide conservation at individual positions and coordinated variation between positions. As is discussed in [2], this MSA is of very limited size and quality, since the 817 aligned sequences are frequently highly gapped, and any generative model trained on these data is expected to be of limited accuracy and generative capacity. Consequently, Lambert et al. [2] did not freely generate sequences, but variants at several mutational distances from an experimentally tested reference wildtype.

DCA represents the family distribution using a Potts model. For a sequence *x* = (*x*_1_, …, *x*_*L*_), the model assigns the generative energy

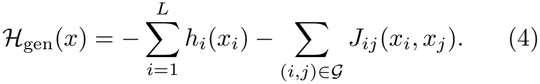

The position-specific fields (or biases) *h*_*i*_(*x*_*i*_) describe the preference for nucleotide *x*_*i*_ at position *i*. They therefore capture conservation patterns, such as nucleotides that are strongly preferred at catalytic or structurally constrained positions. A sequence that introduces an un-common nucleotide at a highly conserved position generally receives an unfavorable field contribution, whereas substitutions at variable positions incur smaller energetic penalties.

The couplings *J*_*ij*_(*x*_*i*_, *x*_*j*_) describe how the contribution of a nucleotide at position *i* depends on the nucleotide found at position *j* on the interaction graph *G*. Each *J*_*ij*_ is a 5 × 5 matrix containing a distinct contribution for every pair of nucleotide or gap states. Consequently, a substitution that is unfavorable in one sequence con-text can become favorable when accompanied by a compensatory substitution at another position. In RNA, such covariation frequently arises from the maintenance of base-pairing interactions: for example, disruption of a Watson–Crick pair by a mutation on one strand can be compensated by a second mutation on the opposite strand. The couplings can also capture constraints associated with tertiary contacts or other forms of epistasis, although statistical coupling does not by itself establish a direct physical interaction.

The parameters *h* and *J* are inferred from the MSA so that the model reproduces the observed single-position frequencies and pairwise correlations. The resulting sequence distribution is analogous to Eq. (2), where *ℋ*_gen_ is the DCA Hamiltonian (4).

Across diverse RNA and protein families, functional sequences have been observed to preferentially occupy low-DCA-energy regions of sequence space. A favorable DCA score therefore indicates compatibility with the evolutionary statistics of the family, which appears to be necessary for function but is not sufficient. DCA measures compatibility with the evolutionary statistics of the family; it cannot guarantee activity under the specific conditions imposed by a given experiment. Indeed, even naturally occurring family members may show little or no detectable activity when tested outside their native molecular or environmental context.

In the group I intron setting, sequences favored by DCA show particularly strong agreement with experimental functionality. Lambert et al. [2] found that approximately 60% of DCA-generated variants remained active at a Hamming distance of 30 substitutions from the *Azoarcus* wild type, whereas randomly mutated sequences showed no detectable activity even at a distance above 10. Detectable activity among DCA designs extended to approximately 60 substitutions, i.e. to slightly more than 30% of the full-lenth wildtype sequence. We therefore use the DCA energy *ℋ*_gen_ from the empirically validated model of Lambert et al. [2] as the plausibility term in the effective Hamiltonian.

### D. A physical score for thermal robustness

The second component of the effective Hamiltonian must provide a sequence-dependent signal for thermal robustness without relying on labeled variants from the target family. We derive this signal from the Turner nearest-neighbor model of RNA secondary-structure thermody-namics [11]. The model assigns a folding free energy Δ*G*_*T*_ (*s* |*x*) to a sequence *x* adopting secondary structure *s* at temperature *T*, by decomposing the structure into contributions from base-pair stacks, loops, and other local motifs. Because these parameters describe interactions determined by nucleotide chemistry rather than properties learned from a particular RNA family, the same model can be applied to any RNA sequence without family-specific training or experimental feedback.

Functional group I introns must adopt a conserved secondary structure that organizes the catalytic core and its supporting helices. Let *C*_Azo_ denote the set of base pairs in the consensus secondary structure of the *Azoarcus* intron [5]. A natural first strategy would be to favor sequences with a low folding free energy Δ*G*_*T*_ (*C*_Azo_ |*x*) for this target structure. The free energy of the target structure alone, however, does not account for folding stability against alternative folds. A mutation can stabilize the consensus structure while stabilizing an incompatible structure even more strongly. Consequently, the absolute free energy of the target fold provides an incomplete measure of whether the sequence preferentially adopts the functional structure.

We therefore work with the equilibrium base-pairing probabilities of the full Turner ensemble. For a sequence *x* at temperature *T*, the probability that positions *i* and *j* are paired is

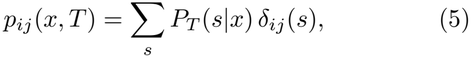

where *δ*_*ij*_(*s*) = 1 if *i* and *j* are paired in *s* and 0 otherwise. *P*_*T*_ (*s*|*x*) *∝*exp[−Δ*G*_*T*_ (*s* |*x*)/*RT*] is the Boltzmann weight of structure *s*, where *R* is the universal gas constant. Then, we define the structure-based (SB) potential as

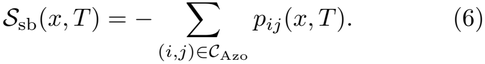

Writing

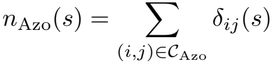

for the number of *Azoarcus* consensus contacts formed in any structure *s*, the quantity *−S*_sb_ is the expected value of *n*_Azo_ under the Turner distribution over secondary-structure space. Low values of *S*_sb_ correspond to ensembles in which more of the consensus structure contacts are formed in the thermodynamic equilibrium of secondary structure folds.

This ensemble-based quantity differs fundamentally from the free energy of a single target structure. Each *p*_*ij*_ is computed relative to all competing structures in the partition function, and each nucleotide can pair with at most one partner in any given structure. Increasing the probability of a consensus contact therefore shifts probability away from structures in which that contact is absent or replaced by an alternative pairing. The potential consequently rewards not only energetic stabilization of the target contacts, but also their specificity relative to competing folds.

Analyzing the Lambert et al. [2] dataset, we find that *S*_sb_(*x*, 37 °C) separates variants that remain active at 60 °C from variants that splice at 37 °C but lose activity under thermal stress (Fig. 2). This separation is stronger than that obtained from the folding free energy Δ*G*_*T*_ (*C*_Azo_ |*x*) of the consensus structure alone (Fig. S3 and Sec. S3.5). We therefore use *S*_sb_ as our physical steering potential.

**FIG 2.**
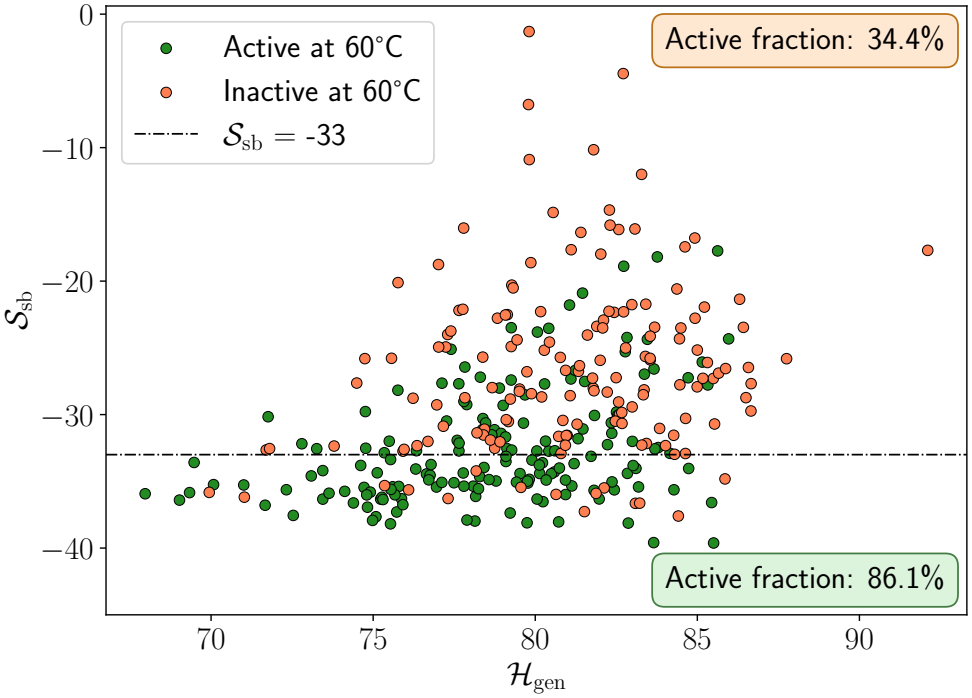
Generative energy ℋ_gen_ and structure-based potential S_sb_ for variants from Lambert et al. [2] that are active at 37 ^°^C and lie between 25 and 30 substitutions from wild-type Azoarcus. Green and orange points denote variants classified as active and inactive, respectively. The horizontal line marks S_sb_ = −33; the active fraction is 86.1% below this cutoff and 34.4% above it. Analysis based on data from Lambert et al. [2]

Secondary-structure stability alone is a good signal for thermal robustness, but it does not imply functionality. Lambert et al. [2] found that generating *Azoarcus* wild-type variants by optimizing on this criterion alone, without an evolutionary constraint, produced sequences that had already lost self-splicing activity by roughly 20 substitutions from the wild type. A sequence can maximize the ensemble probability of the consensus base pairs while violating the sequence constraints imposed by the tertiary contacts and the catalytic core, which are invisible to a secondary-structure model. The two terms therefore address different failure modes, and neither is sufficient on its own.

It must be noted that, unlike the two terms in the MNIST toy case, the DCA and thermodynamic terms are not independent. Variants with both favorable DCA energies and favorable *S*_sb_ values were previously found to be enriched for activity at 37 °C, even though a low *S*_sb_ was not required for activity at that temperature [2]. Part of this dependence is expected: the couplings *J*_*ij*_ are inferred from covariation that largely reflects the maintenance of base pairing, so ℋ_gen_ itself encodes compatibility with the consensus secondary structure. The two terms nevertheless play different roles: ℋ_gen_ keeps the sampled sequences within the evolutionary constraints of a viable group I intron, while *S*_sb_ pushes them toward ensembles in which the consensus contacts form with high probability relative to competing structures.

### E. Sampling the score plane

For each of the ten starting sequences, we sample mutants from the effective Hamiltonian obtained by substituting the two scores defined above into Eq. (3),

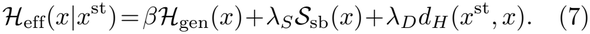

The three Lagrange multipliers control, respectively, how strictly the sampled sequences remain within the evolutionary constraints of the family, how strongly they are driven toward ensembles in which the consensus contacts dominate, and how far they are allowed to move from the starting sequence.

Although the two scores are partially correlated, optimizing either one alone does not ensure favorable values of the other. To separate their contributions experimentally, we therefore sampled three regions of the score plane: the *DCA-favored group*, with favorable evolutionary scores but unfavorable structure-based scores (*↓*ℋ_gen_, *↑S*_sb_); the *SB-favored group*, with favorable structure-based scores but unfavorable evolutionary scores (*↓*ℋ_gen_, *↓S*_sb_); and the *jointly favored group*, with favorable values of both scores (*↓*ℋ_gen_, *↓S*_sb_). Sampling each region generally requires tuning both score weights. All three groups were subject to the same mutational-distance constraint. The remaining region, in which both scores are unfavorable, was not sampled. Fig. 3 shows the three sampled regions of the (ℋ_gen_, *S*_sb_) plane for one of the starting sequences. Region thresholds and numbers of designs are summarized in Table I. The library construction and sampling settings are described in the SI (Sec. S4).

**TABLE I.**
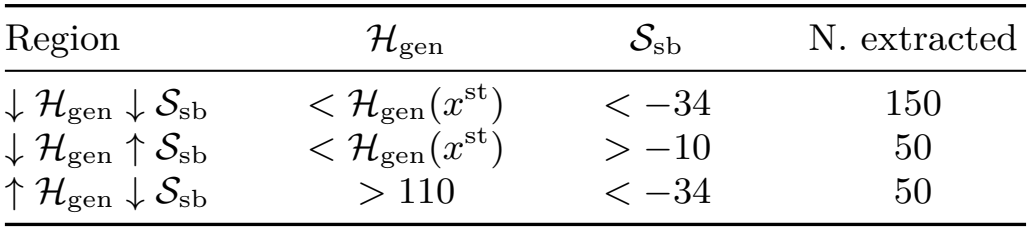
Summary of mutants extracted from different regions of the phase space defined by the DCA energy (ℋ_gen_) and the SB potential (S _sb_). The cutoff S_sb_ < −34 used for the generation is slightly stricter than the value −33 that best separates thermostable from non-thermostable variants in Fig. 2, so as to keep a margin. For further details refer to Sec. S4.4.

| Region | $\mathcal{H}_{\mathrm{gen}}$ | $\mathcal{S}_{\mathrm{sb}}$ | N. extracted |
|---|---|---|---|
| $\downarrow \mathcal{H}_{\mathrm{gen}} \downarrow \mathcal{S}_{\mathrm{sb}}$ | $< \mathcal{H}_{\mathrm{gen}}(x^{\mathrm{st}})$ | < −34 | 150 |
| $\downarrow \mathcal{H}_{\mathrm{gen}} \uparrow \mathcal{S}_{\mathrm{sb}}$ | $< \mathcal{H}_{\mathrm{gen}}(x^{\mathrm{st}})$ | > −10 | 50 |
| $\uparrow \mathcal{H}_{\mathrm{gen}} \downarrow \mathcal{S}_{\mathrm{sb}}$ | > 110 | < −34 | 50 |

**FIG 3.**
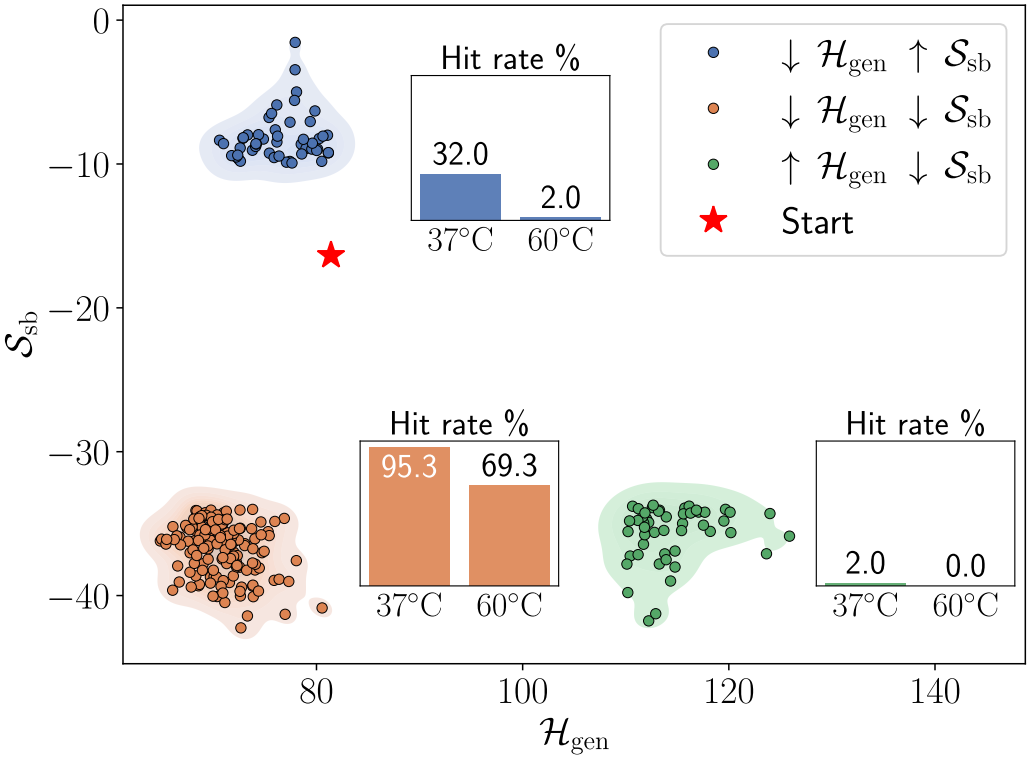
Phase plane obtained for the start sequence n.5. The insets show the hit rates at 37 and 60 ^°^C for the DCA-favored (blue), SB-favored (green), and jointly favored (orange) groups.

To characterize how the designs alter the consensus secondary structure, we calculated its Turner free energy. Since hydrogen bonding is one of the main contributions to this free energy, we also estimated the hydrogen-bond count from the identities of the paired nucleotides to help interpret the calculated energy changes. We report changes in both quantities relative to the corresponding starting sequence.

In total, 250 designs per starting sequence, carrying 5 to 10 substitutions relative to their starting sequence, were synthesized as part of a single oligonucleotide pool and transcribed *in vitro* (see Sec. IV). The designed sequences were incubated with substrates for the self-splicing-like reaction at either 37 or 60 °C. During the first reaction step, the mini-exon, initially attached to the ribozymes 3^*′*^-ends, was transferred to the first substrate, after which the functional ribozymes attached part of the second sub-strate to their 3^*′*^-ends. The raw activity score was calculated as log_10_(*f*_sel_/*f*_ref_), where *f*_sel_ and *f*_ref_ denote each sequence’s frequency in the reacted sample and the control sample not subjected to the reaction, respectively. Two additional groups were included in every pool. A *noise control group* of 1000 sequences with more than 100 substitutions from the *Azoarcus* wild type characterizes the experimental noise background, that is, the signal measured for sequences that are not expected to splice. An *overlap group* of 355 highly-active sequences in the original experiment, common to all pools, is used to place pools measured in separate experiments on a common activity scale. Sampling procedure, region thresholds, selection criteria, reaction conditions, barcoding scheme, and read-processing pipeline are given in Sec. IV and in the SI (Secs. S4, S5 and S6).

### F. Both scores enable self-splicing at 60 °C

Designs in the jointly favored group consistently recover self-splicing at 60 °C, whereas designs in the DCA-favored and SB-favored groups do not. To compare pools assayed in separate experiments, we subtracted the mean log_10_(*f*_sel_/*f*_ref_) of the overlap group from each sequence’s log_10_(*f*_sel_/*f*_ref_), using the same pool and temperature; all activities reported below refer to this normalized loga-rithmic scale. Because the noise distribution measured in the noise control group is heavy-tailed (Fig. S5 and Sec. S6.5), a threshold based on Gaussian assumptions would be inappropriate. We therefore defined the thresh-old empirically, classifying a sequence as a *hit* if its enrichment was at least 10% of the geometric mean en-richment of the overlap group (normalized log_10_ activity *≥ −* 1), and the *hit rate* of a group as the fraction of its sequences classified as hits. As shown in Fig. 4, this cutoff lies above the 99.9th percentile of the noise control sequences activity distribution at both temperatures, making it very unlikely that the activity of a hit arises from measurement noise alone.

**FIG 4.**
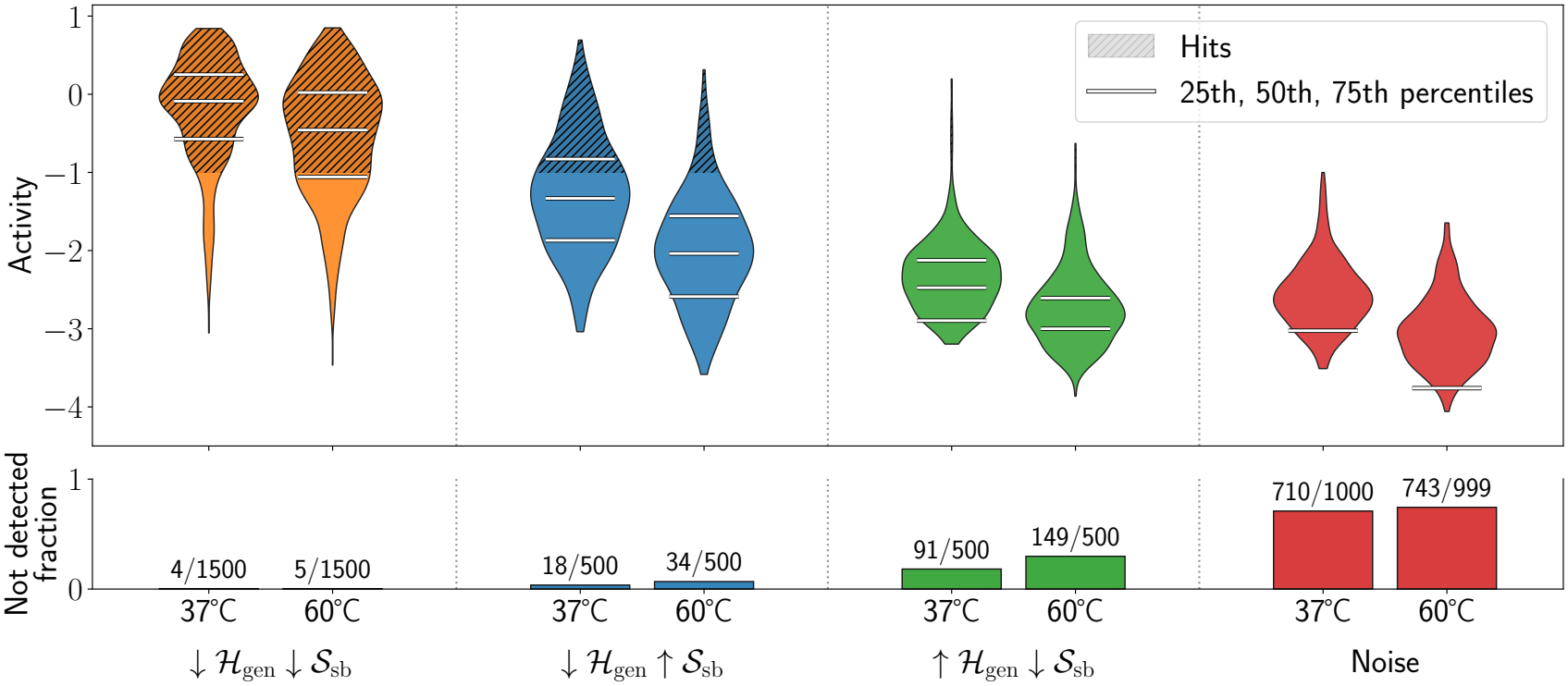
Measured activity for the DCA-favored (blue), SB-favored (green), and jointly favored (orange) groups and the control sequences (noise). For each group—identified by a different color—the two violins represent, from left to right, the activity distributions of the ribozymes measured at 37 ^°^C and 60 ^°^C. The bottom panel shows the fraction of sequences with undetected activity (f_ref_ > 0, f_sel_ = 0), assigned −∞ when computing quantiles. Sequences with f_ref_ = 0 are treated as missing data.

Fig. 3 shows the hit rates at 37 °C and 60 °C for the three groups derived from one starting sequence. The jointly favored group has high hit rates at both temperatures. The DCA-favored group contains active variants at 37 °C, but few remain hits at 60 °C. The SB-favored group contains almost no hits at either temperature. The same pattern holds across starting sequences (Sec. S7, Table S5 and Fig. S8).

Results pooled across starting sequences are shown in Fig. 4 and reported in Table II. The jointly favored group reaches hit rates of 88.9% at 37 °C and 72.5% at 60 °C, compared with 32.2% and 7.8% for the DCA-favored group and 1.0% and 0.4% for the SB-favored group. These comparisons suggest distinct roles for the two scores: favorable DCA energy is strongly associated with activity at 37 °C, whereas favorable base-pairing en-sembles help preserve activity at 60 °C when combined with favorable DCA energy.

**TABLE II.**
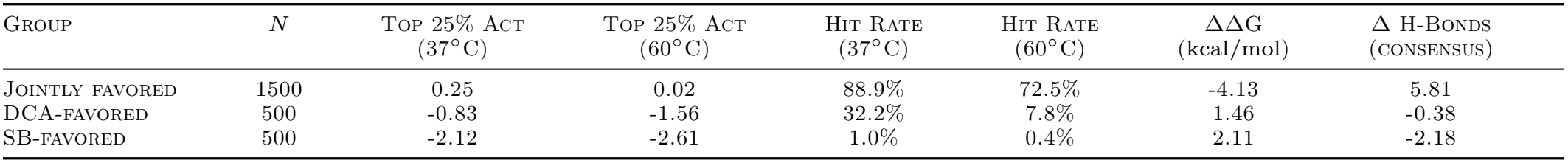
Activity and structural descriptors pooled across the ten starting sequences for the DCA-favored, SB-favored, and jointly favored groups. “Top 25% Act” is the reference-normalized activity delimiting the highest-activity quartile (the 75th percentile), and the hit rate is the fraction of designs with reference-normalized activity in log of at least −1. ΔΔG = ΔG_design_ −ΔG_st_ is the mean change in the Turner free energy of the consensus secondary structure (without the pseudoknot), in kcal/mol, and Δ H-Bonds is the mean change in its number of hydrogen bonds, both relative to the corresponding starting sequence. Designs with an undetected activity (f_sel_ = 0 with f_ref_ > 0) are kept at −∞ in the quantiles and in the hit rates, as in Fig. 4 and Sec. S6.3; sequences with f_ref_ = 0 are treated as missing data. The breakdown by starting sequence, with the numbers of undetected designs, is given in Table S5.

The SB-favored group is the least active of the three (Fig. 4, green), consistent with the absence of functional variants above a DCA energy of 110 reported by Lambert et al. [2]. Its activity distribution is nevertheless shifted above the noise background: its median exceeds the 75th percentile of the noise control distribution, even though nearly all its sequences fall below the hit threshold. The DCA-favored group is more active at 37 °C (blue), but its hit rate drops sharply at 60 °C, mirroring the behavior of the starting sequences. The jointly favored group (orange) combines high activity at 37 °C with substantial retention of activity at 60 °C.

The structural descriptors provide a further indication of why the SB potential is insufficient on its own (Table II). In the SB-favored group, a favorable SB score coincides with an increase in the calculated free energy of the consensus structure (ΔΔ*G* = +2.11 kcal/mol) with a loss of approximately two hydrogen bonds. Both changes run counter to a simple expectation that a favorable SB score should accompany stabilization of the consensus structure. In the jointly favored group, the corresponding changes are −4.13 kcal/mol and approximately six additional hydrogen bonds, consistent with that expectation. As shown above, consensus-structure free energy did not distinguish high-temperature active from inactive variants (Fig. S3), so it should not be treated as a measure of thermal robustness in this dataset. Even so, the unfavorable structural changes and poor activity in the SB-favored group suggest that favoring the SB score while allowing unfavorable DCA scores can favor directions in sequence space that do not preserve other structural features or catalytic activity.

The specific contribution of *S*_sb_ to thermal robustness can be isolated from the change in activity between temperatures. For each sequence, we computed Δ = *a*_60_ − *a*_37_, where *a*_*T*_ denotes the overlap-normalized log_10_ activity at temperature *T*. More negative values of Δ indicate greater activity loss upon heating, while values closer to zero indicate greater activity retention. We restricted this comparison to sequences that exceeded the hit threshold at 37 °C, thereby excluding variants that were already non-functional and separating thermal ro-bustness from baseline activity.

The distributions of Δ shown in Fig. 5 highlight the effect of optimizing *S*_sb_: relative to the DCA-favored group, the distribution for the jointly favored group is shifted by 0.6 decades toward higher values, corresponding to approximately fourfold greater relative activity retention upon heating. Because the two groups share the same starting sequences and mutational budget and have comparable DCA energies, the difference in retained hit rate and the shift in Δ can be attributed to the secondary-structure term.

**FIG 5.**
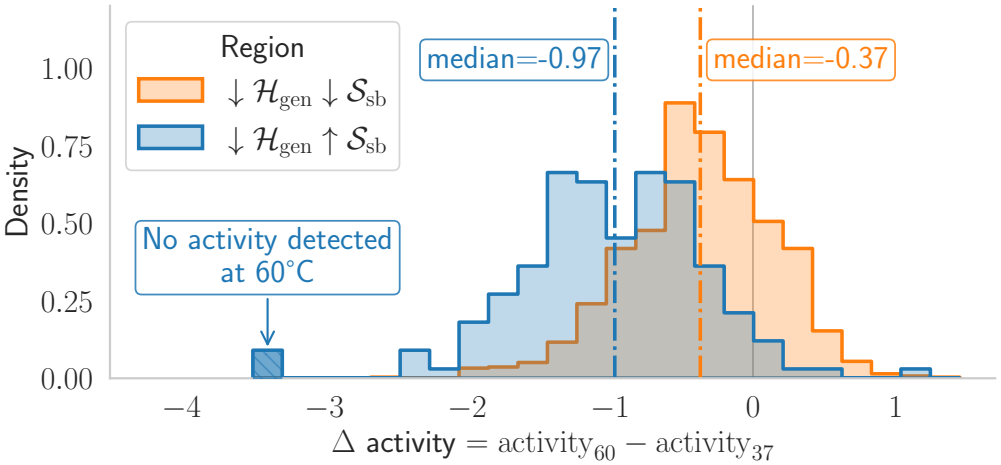
Histograms of the activity change Δ = a_60_ − a_37_, with more negative values indicating greater activity loss, for the DCA-favored and jointly favored groups. Only the sequences with activity above the hit threshold at 37 ^°^C were considered. Sequences with undetected activity at 60 ^°^C (a_60_ = −∞) are placed in the bin at −3.5 for display only.

Overall, the high-throughput experiment validated high-temperature function recovery with only 5 to 10 substitutions—2.5% to 5% of the sequence—and without measurements of thermal stability in the target family.

### G. Thermal robustness and catalytic pocket organization

The limited number of substitutions introduced by our design strategy allows us to investigate how mutations affect thermal robustness within a shared sequence back-ground. We focused on Start–10 and two variants with contrasting temperature responses, selected from the jointly favored (robust) and DCA-favored (sensitive) groups. Both carry nine substitutions relative to Start– 10 and have comparable DCA energies (76.7 and 73.9), but different structure-based scores (−36.2 and −9.6). While both are active at 37 °C, their responses to heating differ markedly: the robust variant increases its normalized log_10_ activity by 1.02, whereas the sensitive variant decreases by 1.53.

We performed explicit-solvent molecular dynamics simulations of wild-type *Azoarcus* (WT), Start–10, and the two variants at 37 and 60 °C, starting from sequence-specific AlphaFold 3 models (Sec. IV E). All systems remained broadly compact over the analyzed 0–300 ns interval, consistent with the persistence of a shared fold on the simulated timescale. Nevertheless, the robust variant exhibited smaller per-nucleotide displacements across much of the sequence at 60 °C, including at the terminal catalytic guanosine ΩG (Sec. S8.4 and Fig. S9). This motivated a closer examination of the catalytic pocket.

We defined the pocket *∂*ΩG as ΩG together with the nine simulated nucleotides with a heavy atom within 5 Å of any ΩG heavy atom in the experimental structure 9V7P [5] (Fig. 6A,B). To quantify its organization, we compared the distances between nucleotide base centroids for all 45 unique pairs in each simulated frame with their reference values in 9V7P. The resulting pocket distance RMSD measures the overall deviation of these internal distances from the experimental geometry, independently of global translation and rotation (Sec. S8.6). The cumulative distributions (Fig. 6E,F) report the percentage of frames with a pocket RMSD below each threshold. They show that the robust variant samples pocket geometries closer to the experimental reference than Start–10 or the sensitive variant. This separation is modest at 37 °C but becomes more pronounced at 60 °C, where Start–10 and the sensitive variant show largely overlapping distributions. WT remains closest to the reference at both temperatures. Segment and time-window sensitivity analyses are reported in Sec. S8.7.

**FIG 6.**
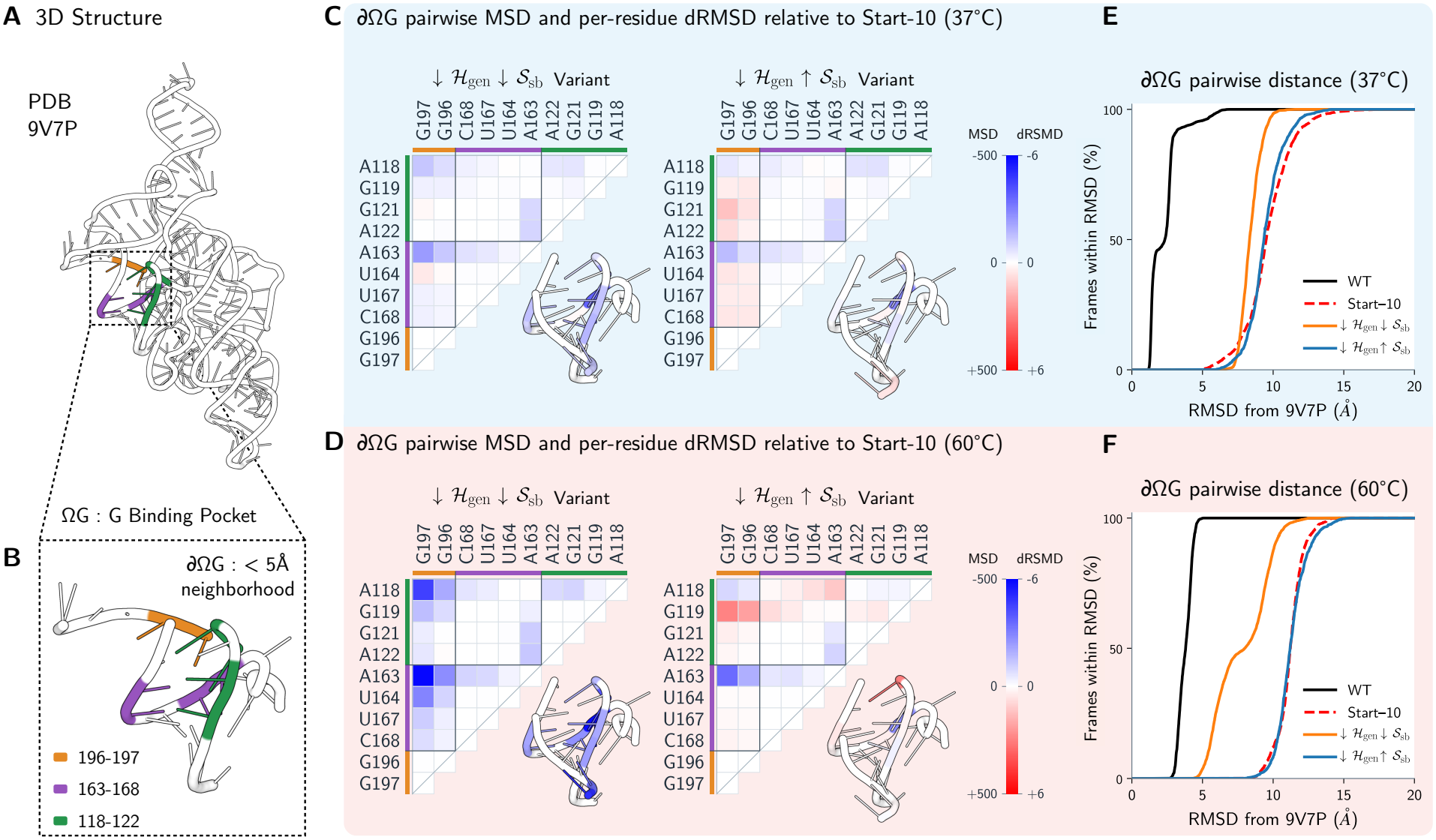
Pocket organization in variants with contrasting thermal responses. (A,B) Experimental structure PDB 9V7P and the fixed ten-nucleotide selection ∂ΩG surrounding terminal guanosine ΩG. (C,D) Pairwise mean-squared deviation (MSD) maps of distances from the reference at 37 and 60 ^°^C for the robust and sensitive variants relative to Start–10. Each cell gives ΔMSD_ij_ (Sec. S8.6), with blue indicating a smaller mean-squared deviation from the reference than Start–10, and red a larger deviation. Structural insets show the corresponding per-residue distance-RMSD differences, in Å (Sec. S8.6). (E,F) Cumulative percentages of frames whose pocket distance RMSD from 9V7P is at or below the indicated threshold at 37 and 60 ^°^C. Pocket distance RMSD is computed over all 45 unique pairs within the fixed selection. Curves show WT (black), Start–10 (dashed red), the robust variant (orange), and the sensitive variant (blue). All analyses use the common 0–300 ns interval.

Pairwise maps further reveal how these differences are distributed within the pocket (Fig. 6C,D). For each nucleotide pair, they report the difference between a variant and Start–10 in the time-averaged squared deviation of their centroid distance from the experimental reference. Negative values (blue) indicate smaller deviations than Start–10, whereas positive values (red) indicate larger deviations. At 60 °C, the robust variant exhibits broadly smaller deviations, including for pairs involving ΩG, whereas the sensitive variant shows a mixture of improvements and larger deviations. These differences primarily reflect changes in mean pocket geometry rather than fluctuations around it (Sec. S8.8).

Importantly, neither variant contains substitutions within *∂*ΩG, suggesting that mutations elsewhere influence the organization of this functional region. A possible contribution involves the P5/P5a receptor, which supports P9 tetraloop docking [25]. Both variants carry C71U, potentially replacing an A–C mismatch with an A–U pair, whereas the sensitive variant additionally carries C59A and G68A, replacing a potential C–G pair with an A–A mismatch. Such changes could modify local pairing and helix packing, indirectly affecting the catalytic pocket, consistent with cooperative coupling between distant RNA structural elements [6, 26].

Together, these simulations associate retained high-temperature activity with reduced whole-chain displacements and a more reference-like organization of an un-mutated catalytic pocket. They suggest that a limited number of substitutions can influence distant functional regions through changes in the broader RNA architecture.

## III. CONCLUSIONS

In this work, we have shown that biophysical information can complement evolutionary sequence models to guide the design of functional RNAs toward improved thermal robustness. By combining a DCA generative model with a structure-based potential derived from RNA secondary-structure thermodynamics, we recovered high-temperature self-splicing activity in ribozymes previously inactive at 60 °C, using only 5–10 substitutions and without training on experimental measurements of the target property.

Our results highlight the complementary roles of the two sources of information. Evolutionary constraints favor sequences compatible with the functional RNA family, while the physical potential favors sequences associated with greater thermal robustness. The underlying thermo-dynamic parameters describe interactions such as base stacking and pairing, which are common to all RNA molecules and do not require family-specific retraining. However, this physical description alone does not capture all the constraints necessary for catalytic function. Neither the evolutionary nor the structure-based criterion was sufficient on its own, whereas their combination yielded a substantial fraction of active variants at 60 °C. More broadly, our findings illustrate a general strategy for property-directed molecular generation based on combining evolutionary sequence models with complementary sources of information. Within the effective-Hamiltonian framework, the steering potential could be provided by an auxiliary predictor trained on experimental measurements when sufficient data are available, or by a physical model constructed independently of the target family, as demonstrated here. The latter is particularly valuable in data-scarce settings, where learning an accurate property predictor may be impractical. Rather than necessarily requiring larger experimental datasets, this approach seeks to make better use of existing knowledge by integrating complementary sources of information to guide sequence design.

## IV. METHODS

### A CA training

Sequence plausibility was quantified using the Potts model Hamiltonian (4) on a sparse graph *G*. We used the multiple sequence alignment and pretrained edge-activation DCA model from Lambert et al. [2]. The training procedure is described in [13] and implemented in the Python package `adabmDCA 2.0` [12] (`--model edgeDCA` in the training script). Briefly, instead of tuning the couplings between every pair or residues in a fully-connected fashion, edge-activation DCA starts from an empty interaction graph (*J*_*ij*_(*·,·*) = 0 *∀ i, j* = 1, …, *L*). At each iteration, it selects the pair of residues (*i, j*) whose coupling yields the largest reduction in the Kullback-Leibler divergence between the current model and the data distribution, and adds the corresponding edge *J*_*ij*_(*·,·*) *∈* ℝ ^5×5^ to the graph. To construct the training alignment, a seed containing the *Azoarcus* sequence and the secondary structure derived from PDB 1U6B [27] was prepared. Then, a homology search was done using Infernal in the Rfam RF00028 family with the E-value threshold set at 10^−3^. The final alignment contains 817 sequences of length *L* = 193, corresponding to the 197-nucleotide construct without the first three nucleotides (GUG) and the terminal G. DCA positions 1–193 therefore correspond to construct positions 4–196; residue numbering elsewhere refers to the full construct. The DCA model was trained on this dataset until the Pearson coefficient between the natural sequences and the model connected correlations was 0.95. The final sparse graph contains ~ 12% of the parameters of a fully-connected model. The variants generated by this model and tested in Lambert et al. [2] (among which there are the 10 starting sequences that we considered) were produced by mutating the *Azoarcus* sequence using only substitution of nucleotides. In particular, the emission of a gap symbol was not permitted. For a complete description of the model, training alignment, software settings, and sequence-position mapping, see SI Sec. S3, particularly Secs. S3.1 and S3.2.

### B. Structure-based score

The structure-based potential, used as the thermal stability phenotype, was calculated from the equilibrium secondary-structure ensemble of each RNA sequence using ViennaRNA [28]. Pairing probabilities *p*_*ij*_(*x, T*) were obtained by partition-function calculations under the thermodynamic model, without imposing the reference structure as a folding constraint. The reference dot-bracket structure was converted into a binary contact map *δ*_*ij*_(*C*_Azo_), identifying its prescribed base pairs. The physical potential was then defined as *S*_sb_(*x, T*) = − ∑_*i*_ *δ*_*ij*_ (*C*_Azo_)*p*_*ij*_ (*x, T*), i.e., the negative sum of the ensemble probabilities of the reference contacts, without normalization by sequence length or contact number. All SB scores used for sequence design and analysis were calculated at *T* = 37 °C; analogous results were obtained at *T* = 60 °C (see Secs. S3.3, S3.4 and S3.6, Figs. S2 and S4). More negative values indicate greater thermo-dynamic support for the reference pairing pattern. This dimensionless potential measures structural compatibility with the reference rather than a folding free energy.

### C. Multi-step Monte Carlo sampling

Computing the physical potential using ViennaRNA for a pool of candidate variants is computationally expensive, making an evaluation at each move of a long Monte Carlo simulation almost prohibitive. To sample starting sequence variants using the effective Hamiltonian (7), we therefore implemented a multi-step approach: we propose a sequence of moves (10 in our simulations) using the “fast” terms of the Hamiltonian, consisting of the DCA energy and the Hamming distance penalty, and then we accept or reject the whole set of moves using the “slow” thermodynamic term [29]. The combination of these two steps satisfies detailed balance for the joint Hamiltonian, and the simulation is guaranteed to asymptotically converge to the effective equilibrium distribution defined by In our simulations, “fast” move corresponds to *L* attempted mutations. Each of these attempts selects a residue *i ∈* {1, …, *L}* at random and emits a nucleotide *a* using the conditional Gibbs probability

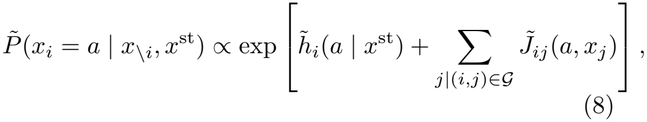

where we defined the modified parameters

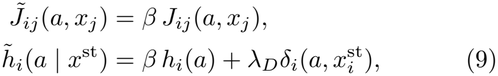

with 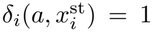 if 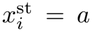 and 0 otherwise. Thus, increasing positive *λ*_*D*_ favors matches to the starting sequence and reduces the mutational distance. Notice that the already present nucleotide can be re-emitted, which corresponds to rejecting the move. Following Lambert et al. [2], the local field corresponding to the gap symbol was artificially set to a very negative value, so that the model never proposed the insertion of a gap. A sequence of 10 “fast” moves transforms an initial state *x*^i^ into a final state *x*^f^, which is accepted with the “slow” move probability

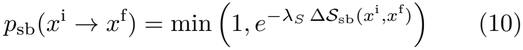

where Δ*S*_sb_(*x*^i^, *x*^f^) =*S*_sb_(*x*^f^) *− S*_sb_(*x*^i^).

In practice, our simulations consist of 200 “fast” steps and 20 “slow” steps in total, which were checked to bring the sampling close to the effective model’s equilibrium. We tuned *β* and *λ*_*S*_ in order to target the desired region of the (ℋ_gen_, *−S*_sb_) plane, and chose a value *λ*_*D*_ for which the bulk of the histogram of distances from the starting sequence was between 5 and 10 substitutions. The pool of variants for each group and wild type was filtered using the criteria of Tab. I, plus the requirement of being 5 to 10 mutations away from the starting sequence (see SI Sec. S4.4 for distance constraints relative to *Azoarcus*). In this way, we avoid the trivial result that the variants become more thermostable than the starting sequences because they moved towards the wild type. The detailed-balance derivation, the sampling parameters for each group, the tuning procedure for the distance penalty, the distance constraints and the filtering of the library are given in Secs. S4.1–S4.4 and Table S2.

### D. High-throughput activity screening

Designed ribozymes were screened using a pooled self-splicing-like assay previously validated on libraries of up to 18,000 molecules [2, 13]. RNA pools were produced by PCR amplification and *in vitro* transcription of synthe-sized DNA templates, followed by purification. Reactions containing 2 *µ*M RNA and 25 *µ*M each of substrates S1 and B-S2 were incubated for one hour at 37 or 60 °C in 30 mM EPPS (pH 7.5) and 60 mM MgCl_2_, then stopped with EDTA. A zero-time RNA control was processed using the same purification procedure.

Reacted and control samples were converted into sequencing libraries and analyzed by Oxford Nanopore sequencing. Reads were assigned to designed sequences to estimate their frequencies in the control pool, *f*_ref_. Reacted reads were identified by removal of the 3^*′*^ exon and attachment of substrate S2, yielding the selected frequencies *f*_sel_. Experimental protocols, construct and substrate sequences, library preparation, and read-assignment details are provided in the SI (Secs. S5 and S6, Table S3). Sequencing read counts and coverage are reported in SI Table S4.

Raw log activity was defined as *b* = log_10_(*f*_sel_/*f*_ref_). The reference set comprised the 355 overlap-group sequences included in every pool (reference-group definition and normalisation in Sec. S6.4). Their mean log activity in a given pool and at temperature *T* was denoted 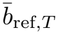. Each design’s normalized activity was its raw log activity minus this reference mean from the same pool and temperature 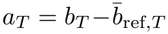. The reference-group properties and the agreement of its measured activities with the previous experiment are shown in SI Fig. S7.

Sequences with *f*_ref_ = 0 were treated as missing data and received no activity score. For *f*_ref_ *>* 0 but *f*_sel_ = 0, activity was classified as undetected and assigned *a*_*T*_ = *−∞*, including when computing quantiles (Fig. 4). The finite bin used for undetected activity in Fig. 5 is a display convention only.

### E. Molecular dynamics simulations

We performed explicit-solvent molecular dynamics simulations of the wild-type *Azoarcus* group I intron, Start– 10, and two Start–10-derived variants with contrasting thermal responses. Initial structures were obtained using AlphaFold 3 [30]. Simulations were performed using GROMACS 2025.1 [31] at 310 and 333 K (approximately 37 and 60 °C). Comparisons used a 0–300 ns analysis window, with sampling, trajectory handling, and averaging detailed in Sec. S8.3 and Table S6. Structure prediction, system preparation [32–34], simulation parameters [35– 37], and trajectory processing [38–42] are detailed in the SI (Secs. S8.2 and S8.3).

Sequence-wide dynamics were characterized using pernucleotide root-mean-square displacements between globally fitted frames separated by 10 ns. To characterize the organization of the catalytic pocket, we defined a fixed selection, *∂*ΩG, comprising terminal guanosine ΩG and the nucleotides represented in the simulations with a heavy atom within 5 Å of ΩG in PDB 9V7P [5]. Pocket geometry was quantified from deviations of inter-base heavy-atom centroid distances from their experimental values, summarized as pocket distance RMSD and pairwise mean-squared distance deviations from the reference. Definitions, residue mappings, trajectory handling, and sensitivity analyses are provided in the SI (Secs. S8.5–S8.7, Table S7, Fig. S10).

## V. CODE AND DATA AVAILABILITY

The code to reproduce the analyses and figures of this work is available on GitHub at https://github.com/spqb/thermostable-ribozymes. The data, including the files and code for the molecular dynamics analyses, have been deposited in Zenodo (DOI: 10.5281/zen-odo.24418219).

## V. ACKNOWLEDGMENTS AND FUNDING

This work was supported by the AISSAI Center (CNRS), funded by the CNRS Foundation through a grant from Google, and by the Horizon Europe MSCA Staff Exchange project “SIMBAD” (grant agreement no. 101131463). This work was also supported by ERC Abio-Evo (101002075) and PEPR Origins (ANR-22-EXOR-0013). We thank Kevin Ricard and the PEPR Molecu-larXiv sequencing platform at UMR Gulliver 7083 (grant ANR-22-PEXM-0002).

