## Supplementary Information for "Biophysics-informed generative models improve activity and thermostability of designed ribozymes"

Francesco Calvanese†,[1,2] Lorenzo Rosset†,[3,2] Polina Pavlinova,[4] Louis Meuret,[5] Vaitea Opuu,[4,6] Samuela Pasquali,[5] Philippe Nghe,[4,6] Francesco Zamponi,[7] and Jorge Fernandez-de-Cossio-Diaz*,[1], Martin Weigt‡,[2,8]

[1] *Institut de Physique Théorique, Université Paris-Saclay, CNRS, CEA, Gif-sur-Yvette, France*
[2] *Sorbonne Université, CNRS, Dept. of Computational, Quantitative and Synthetic Biology - CQSB, Paris, France*
[3] *École Normale Supérieure, CNRS, Laboratoire de Physique - LPENS, Paris, France*
[4] *Laboratoire de Biophysique et Évolution, Chimie Biologie Innovation (CBI), UMR 8231, ESPCI Paris, Université PSL, CNRS, 75005 Paris, France*
[5] *Laboratoire de Biologie Fonctionnelle et Adaptative, Université Paris Cité, CNRS UMR 8251, INSERM ERL 1133, F-75013 Paris, France*
[6] *Laboratoire de Biologie Structurale de la Cellule (BIOC), CNRS, École polytechnique, Institut Polytechnique de Paris, 91128 Palaiseau, France*
[7] *Dipartimento di Fisica, Sapienza Università di Roma, Rome, Italy*
[8] *Institut Universitaire de France (IUF), France*

(Dated: November 21, 2026)

### CONTENTS

S1. Bayesian interpretation of classifier steering 2

S2. Starting sequences 3
S2.1. Selection criteria 3
S2.2. Identity and properties of the ten starting sequences 3
S2.3. Activity versus distance in the previous dataset 3

S3. Model and thermodynamic-score implementation 4
S3.1. Generative model and training alignment 4
S3.2. Mapping between modelled and construct positions 5
S3.3. Consensus structure and contact set 5
S3.4. Thermodynamic calculations 6
S3.5. Consensus free energy as an alternative score 7
S3.6. Structure-based potential at 37 and 60°C 7

S4. Sampling and library construction 8
S4.1. Detailed balance of the multi-step scheme 8
S4.2. Sampling schedule and parameters 10
S4.3. Distance constraints 11
S4.4. Filtering and selection of the library 11

S5. Experimental procedures 11
S5.1. Library composition 11
S5.2. RNA preparation 12
S5.3. Self-splicing-like reaction and control 12
S5.4. Library preparation and sequencing 12

S6. Activity processing and controls 12
S6.1. Basecalling and read assignment 13

† These authors contributed equally to this work.
*
‡

S6.2. Read counts and coverage 13
S6.3. Missing and undetected activities 13
S6.4. Reference group and normalisation 14
S6.5. Noise controls and the hit threshold 15
S6.6. The activity cutoffs of the previous dataset 16
S6.7. Agreement with the previous experiment 17

S7. Results by starting sequence 17
S7.1. Quantities reported 17
S7.2. The ordering of the three regions holds for every starting sequence 18
S7.3. Variability between starting sequences 18
S7.4. The structure-only region 18

S8. MD methods and robustness checks 21
S8.1. Sequences and structure prediction 21
S8.2. System preparation and simulation conditions 21
S8.3. Trajectory processing 22
S8.4. Whole-chain and per-nucleotide displacements 23
S8.5. Pocket selection and residue mapping 24
S8.6. Pocket-distance metrics and structural insets 25
S8.7. Segment and time-window sensitivity 26
S8.8. Mean-geometry shifts and fluctuations 26
S8.9. Displacements outside mutation neighbourhoods 27

References 28

### S1. BAYESIAN INTERPRETATION OF CLASSIFIER STEERING

To make the Bayesian interpretation concrete, we consider the MNIST example from the main text, where $x$ is a binary image and the target label is $y = 8$. Throughout this section, we set $\lambda_D = 0$, so that no constraint on the distance from a starting configuration is imposed.
The generative model defines a prior probability distribution over configurations,

$$P_{\text{gen}}(x) = \frac{e^{-\mathcal{H}_{\text{gen}}(x)}}{Z}, \tag{S1}$$

and the classifier provides the likelihood $P(y = 8 \mid x)$ of the target label given an image. The steering potential is

$$\mathcal{S}_{\text{class}}(x) = -\log P(y = 8 \mid x). \tag{S2}$$

For $\beta = \lambda_S = 1$, the effective Hamiltonian therefore gives the sampled distribution

$$P(x) = \frac{e^{-\mathcal{H}_{\text{gen}}(x) - \mathcal{S}_{\text{class}}(x)}}{\sum_{x'} e^{-\mathcal{H}_{\text{gen}}(x') - \mathcal{S}_{\text{class}}(x')}}. \tag{S3}$$

Using the definitions above, this can be rewritten as

$$\begin{aligned} P(x) &= \frac{P_{\text{gen}}(x) P(y = 8 \mid x)}{\sum_{x'} P_{\text{gen}}(x') P(y = 8 \mid x')} \\ &= \frac{P(x, y = 8)}{P(y = 8)} = P(x \mid y = 8), \end{aligned} \tag{S4}$$

where the joint model is defined by $P(x, y) = P_{\text{gen}}(x) P(y \mid x)$. The factor $1/Z$ from the generative distribution cancels between numerator and denominator, and the remaining normalization is the evidence,

$$P(y = 8) = \sum_{x'} P_{\text{gen}}(x') P(y = 8 \mid x'). \tag{S5}$$

Sampling with the sum of the generative energy and the classifier score can thus be interpreted as conditional generation at the target label $y = 8$. Neither $Z$ nor $P(y = 8)$ needs to be evaluated: Monte Carlo sampling only requires changes in the effective Hamiltonian, computed from the generative energy and the classifier score.
The conditional distribution above corresponds to the choice $\beta = \lambda_S = 1$. More generally, we can raise the prior and likelihood to adjustable powers, a procedure known as tempering, to control their influence on sampling independently. The sampled distribution then becomes

$$\begin{aligned} P_{\beta,\lambda_S}(x) &\propto e^{-\beta\mathcal{H}_{\mathrm{gen}}(x)-\lambda_S\mathcal{S}_{\mathrm{class}}(x)} \\ &\propto P_{\mathrm{gen}}(x)^{\beta} P(y = 8 \mid x)^{\lambda_S}. \end{aligned} \tag{S6}$$

At fixed $\lambda_S$, increasing $\beta$ strengthens the preference for configurations with high probability under the generative model. At fixed $\beta$, increasing $\lambda_S$ strengthens the preference for configurations that the classifier confidently identifies as eights. Adjusting these parameters therefore allows us to steer generation toward the target label while controlling the preference for plausible digits.

### S2. STARTING SEQUENCES

#### S2.1. Selection criteria

All ten starting sequences are taken from the library of *Azoarcus* group I intron variants designed and assayed by Lambert *et al.* [1]. We restricted the choice to the 3429 variants that were assayed at both temperatures, were active at 37 °C ($a_{37} > -2.76$) and had at least eleven reference reads in the 60 °C condition, so that the absence of activity at 60 °C can be read as a measurement rather than as a missing value. Within this subset we required:

- activity at 37 °C above the illustrative cutoff of Ref. [1], $a_{37} > -2.76$, where activities are reported throughout as $a_T = \log_{10}(f_{\mathrm{sel}}/f_{\mathrm{ref}})$;
- no detected activity at 60 °C;
- a Hamming distance from the *Azoarcus* wild type in the range $[25, 30]$;
- a sequence produced by one of the two DCA generative models of Ref. [1] ($T = 1$ and $T = 0.3$), so that the starting sequences lie in the region of sequence space described by the generative model we use here.

The first two requirements select the phenotype we want to repair: a ribozyme that is functional at room temperature but loses its activity at high temperature. The distance window is the regime in which the DCA designs of Ref. [1] are still frequently functional at 37 °C (about 63% of the variants in the 25–30 bin, Fig. S1) while being far enough from the wild type that the residual activity cannot be ascribed to a near-wild-type sequence. It also matches the distance budget imposed on the designs (Table I of the main text), so that the mutants remain in a region of sequence space that has already been probed experimentally.
These criteria leave 445 matched variants in the 25–30 distance bin, of which 126 have no detected activity at 60 °C and 72 come from the two DCA models. From this pool we selected ten sequences spanning the available range of distances and of structure-based scores.

#### S2.2. Identity and properties of the ten starting sequences

Table S1 lists the ten starting sequences with their identifiers in Ref. [1], the generative model that produced them, their distance from the *Azoarcus* wild type, their measured activities and their two scores. Eight starts were generated by the DCA model at $T = 1$ and two by the DCA model at $T = 0.3$.

#### S2.3. Activity versus distance in the previous dataset

Figure S1 shows, for the 7550 variants of Ref. [1] belonging to the three libraries that were assayed at 60 °C in their entirety (DCA at $T = 1$, DCA at $T = 0.3$ and VAE; the remaining libraries were assayed at 60 °C only in part, so that restricting to these three keeps both fractions referred to the same population), variants with an undetected activity being counted as inactive, the fraction that is active at 37 °C as a function of the Hamming distance from the *Azoarcus*

TABLE S1. **Properties of the ten starting sequences.** Identifier and generative model as in Ref. [1]; $d_H$(Azo): Hamming distance from the *Azoarcus* wild type; $a_{37}$, $a_{60}$: activities measured in Ref. [1] ($\log_{10} f_{\rm sel}/f_{\rm ref}$; $-\infty$ = no detected activity); $\mathcal{H}_{\rm gen}$: DCA generative energy; $\mathcal{S}_{\rm sb}(T)$: structure-based potential evaluated at $T = 37$ and 60 °C. The wild type is given for reference; its activities were not measured in the same assay.

| Start | Identifier | Model | $d_H$(Azo) | $a_{37}$ | $a_{60}$ | $\mathcal{H}_{\rm gen}$ | $\mathcal{S}_{\rm sb}$(37 °C) | $\mathcal{S}_{\rm sb}$(60 °C) |
|---|---|---|---|---|---|---|---|---|
| 1 | `dca_f_sequenza_682` | DCA, $T = 1$ | 25 | −1.66 | $-\infty$ | 82.3 | −14.7 | −10.9 |
| 2 | `dca_f_sequenza_689` | DCA, $T = 1$ | 25 | −1.23 | $-\infty$ | 79.8 | −10.9 | −19.2 |
| 3 | `dca_f_sequenza_698` | DCA, $T = 1$ | 25 | −1.87 | $-\infty$ | 83.0 | −21.8 | −17.8 |
| 4 | `dca_f_sequenza_707` | DCA, $T = 1$ | 25 | −1.58 | $-\infty$ | 79.8 | −1.3 | −0.7 |
| 5 | `dca_f_sequenza_739` | DCA, $T = 1$ | 25 | −1.94 | $-\infty$ | 81.4 | −16.4 | −13.2 |
| 6 | `dca_f_sequenza_762` | DCA, $T = 1$ | 30 | −1.51 | $-\infty$ | 77.8 | −22.1 | −23.7 |
| 7 | `dca_f_sequenza_794` | DCA, $T = 1$ | 30 | −2.13 | $-\infty$ | 75.8 | −20.1 | −18.2 |
| 8 | `dca_f_sequenza_799` | DCA, $T = 1$ | 30 | −0.96 | $-\infty$ | 80.6 | −14.9 | −15.2 |
| 9 | `old_dca_619_20_30` | DCA, $T = 0.3$ | 29 | −1.72 | $-\infty$ | 85.2 | −21.9 | −22.8 |
| 10 | `old_dca_953_30_40` | DCA, $T = 0.3$ | 30 | −1.87 | $-\infty$ | 82.3 | −22.3 | −22.2 |
| *Azoarcus* wild type | | — | 0 | — | — | 72.6 | −32.5 | −28.0 |

wild type, together with the fraction of those active variants that also exceed the 60 °C cutoff ($a_{60} > -2/\ln 10 \simeq -0.87$ in $\log_{10}$ units, i.e. −2 in the natural-logarithm units used in Ref. [1]).

Two features of this figure motivate our choice of starting sequences. First, activity at 37 °C decays slowly with distance: 63% of the 606 variants in the 25–30 bin are active, so a starting sequence taken from that bin is a plausible but far from trivial variant. Second, the high-temperature phenotype decays considerably faster: among the variants that are active at 37 °C, the fraction above the 60 °C cutoff drops from 95% in the 0–5 bin to 47% in the 25–30 bin and to 14% and below beyond distance 40, where only a few tens of active variants per bin remain. Thermostability is therefore lost well before room-temperature functionality, and this is precisely the gap that the structure-based steering potential is meant to close.

The asymmetry also holds sequence by sequence: of the 2292 variants of the dataset that are above the 60 °C cutoff, only 20 fall below the 37 °C cutoff, and a single one has no detected activity at 37 °C at all (within the matched subset, none of the 2200 variants above the 60 °C cutoff is inactive at 37 °C). Baseline functionality at 37 °C is thus a prerequisite for high-temperature activity, which is why the ten starts were chosen among variants that already work at 37 °C.

### S3. MODEL AND THERMODYNAMIC-SCORE IMPLEMENTATION

This section records the model files, the position mapping and the software settings needed to recompute $\mathcal{H}_{\rm gen}$ and $\mathcal{S}_{\rm sb}$ for any sequence of the library, and reports the two control comparisons referred to in the main text: the consensus-structure free energy as an alternative score, and the structure-based potential evaluated at 60°C instead of 37°C.

#### S3.1. Generative model and training alignment

The evolutionary score $\mathcal{H}_{\rm gen}$ is the Hamiltonian of the sparse Potts model of Ref. [1], which we use as published, without refitting. The model was obtained with the edge-activation algorithm of Ref. [2] as implemented in `adabmDCA 2.0` [3], trained until the Pearson correlation between the connected two-site correlations of the data and of the model reached 0.95; the resulting interaction graph retains about 12% of the couplings of a fully connected model. Its training alignment was built from a seed containing the *Azoarcus* sequence together with the secondary structure derived from PDB 1U6B [4], followed by an Infernal homology search of the Rfam family RF00028 with an E-value threshold of $10^{-3}$; it contains 817 sequences of length $L = 193$.

Scores are evaluated with the same package and the same conventions as during training. The alphabet is the default RNA alphabet of `adabmDCA 2.0`, `-ACGU`, with the gap symbol in first position, so that the model parameters are a field $h_i(a) \in \mathbb{R}^5$ per position and a coupling $J_{ij}(a, b) \in \mathbb{R}^{5\times5}$ per edge of the graph. The parameter file and the training alignment are provided with the code.

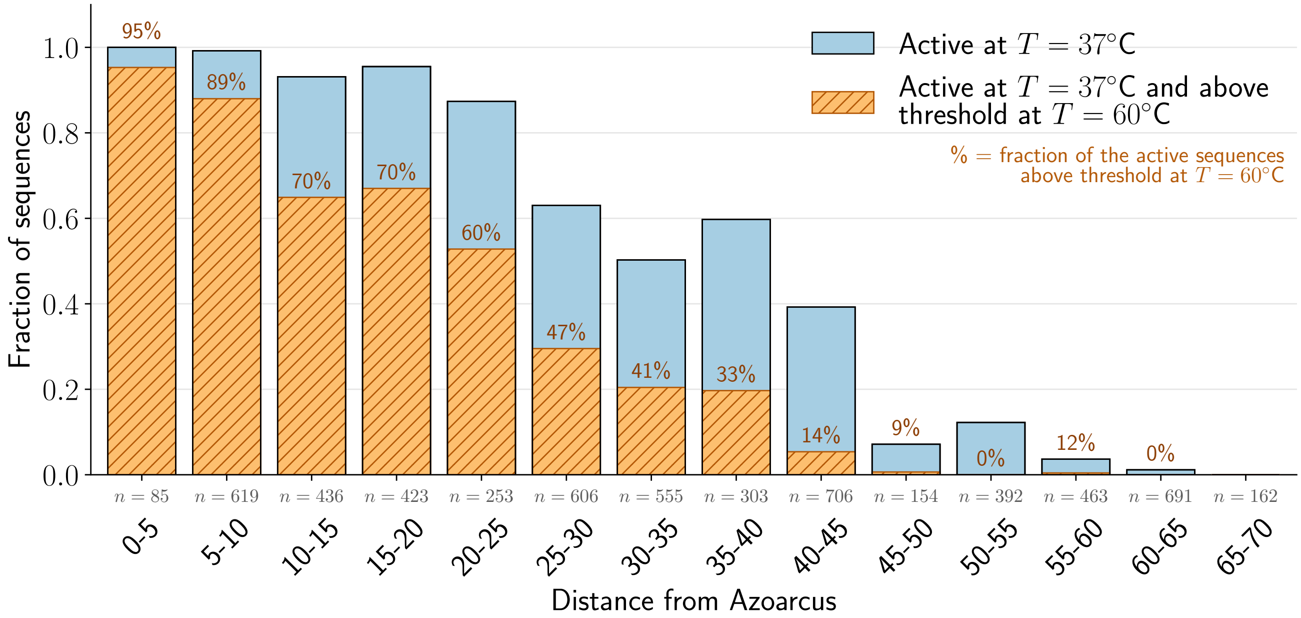


FIG. S1. **Activity versus distance from the wild type in the dataset of Ref. [1].** Blue bars: fraction of the variants of the three libraries assayed at 60 °C in their entirety (DCA at $T = 1$, DCA at $T = 0.3$ and VAE) that are above the 37 °C cutoff ($a_{37} > -2.76$), variants with an undetected activity counting as inactive, in bins of Hamming distance from the *Azoarcus* wild type; $n$ is the number of variants in each bin. Hatched orange: the part of each bar that is also above the 60 °C cutoff ($a_{60} > -2/\ln 10$); the percentage printed above each bar is this fraction relative to the variants active at 37 °C and measured at 60 °C; it is omitted in the last bin, which contains no active variant. The ten starting sequences were selected from the 25–30 bin.

#### S3.2. Mapping between modelled and construct positions

The synthesised constructs are 197 nucleotides long, while the model acts on $L = 193$ positions. The two are related by the constant flanks of the construct,

$$x^{\text{construct}} = \texttt{GUG} \cdot x^{\text{model}} \cdot \texttt{G},$$

so that model positions 1–193 correspond to construct positions 4–196, and construct positions 1–3 and 197 are outside the model. *Azoarcus* carries no gap in the 193 alignment columns, so the correspondence is one-to-one for the wild type and for every variant considered here, all of which were generated by substitution only. Every residue number quoted in this work, in the main text and in this supplement, refers to the full 197-nucleotide construct. Scoring a sequence therefore consists of stripping the four flanking nucleotides, encoding the remaining 193 in the `-ACGU` alphabet and evaluating the Potts Hamiltonian.

The gap symbol is part of the alphabet but is never used by the designs: following Ref. [1], the field of the gap symbol was set to a large negative value during sampling, so that no gap can be emitted, and all designed variants differ from their starting sequence by substitutions only. Gaps therefore play no role in the scores reported here, although they are present in the training alignment and hence in the fitted parameters.

#### S3.3. Consensus structure and contact set

The reference contact set $\mathcal{C}_{\text{Azo}}$ entering $\mathcal{S}_{\text{sb}}(x,T) = -\sum_{(i,j)\in\mathcal{C}_{\text{Azo}}} p_{ij}(x,T)$ is the set of base pairs of the consensus secondary structure of the *Azoarcus* intron derived from PDB 1U6B [4], the same structure used to build the seed alignment. It is stored as a dot-bracket string over the full 197-nucleotide construct and contains 59 base pairs: 53 nested pairs, written with round brackets, and 6 pairs forming the pseudoknot, written with square brackets, which connect positions 119–125 to positions 164–169. Figure S2 shows the structure with the two classes of contacts and with the positions covered by the model.

The contact set is a property of the reference structure alone and is held fixed for every sequence scored: $\delta_{ij}(\mathcal{C}_{\text{Azo}})$ never changes, and only the pairing probabilities $p_{ij}(x,T)$ depend on the sequence. The sum is not normalised by the number of contacts, so $\mathcal{S}_{\text{sb}}$ ranges between 0 and $-59$ in principle; the *Azoarcus* wild type reaches $\mathcal{S}_{\text{sb}} = -32.5$ at 37°C.

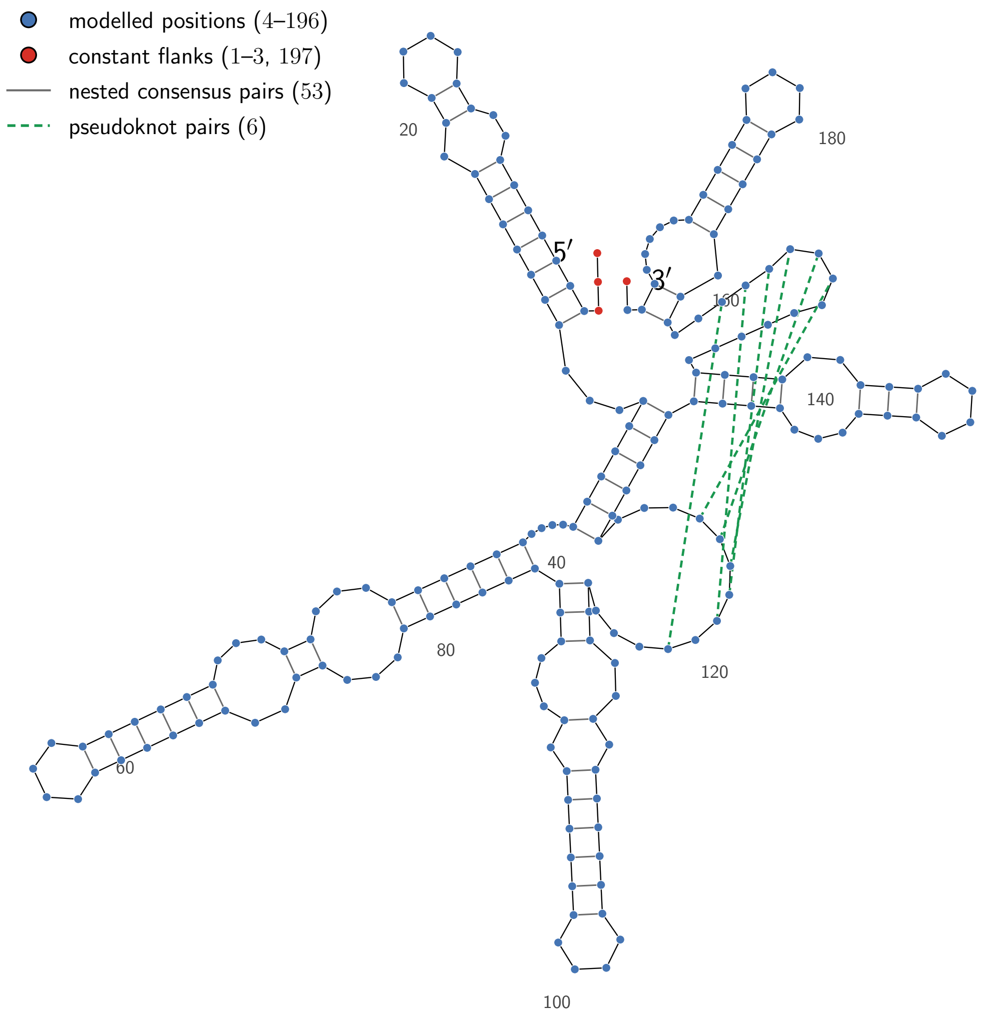


FIG. S2. **Consensus secondary structure of the *Azoarcus* intron, contact set and modelled positions.** The 197-nucleotide construct is drawn with the layout that ViennaRNA assigns to the nested part of the structure; numbers mark every twentieth construct position. Blue circles are the 193 positions covered by the DCA model (construct positions 4–196), red circles the constant flanks `GUG` and `G` that lie outside it. Grey lines are the 53 nested consensus base pairs and green dashed lines the 6 pseudoknot pairs; together they form the contact set $\mathcal{C}_{\text{Azo}}$ summed over in $\mathcal{S}_{\text{sb}}$. The pseudoknot pairs are drawn as chords because a pseudoknot has no planar secondary-structure layout.

#### S3.4. Thermodynamic calculations

All thermodynamic quantities were computed with the Python interface of ViennaRNA [5] version 2.7.2, using the default Turner 2004 nearest-neighbour parameter set. The only model detail changed from its default value is the folding temperature, which is set to 37°C or to 60°C as indicated; dangling-end treatment, the special hairpin tables and the maximum base-pair span are left at their defaults. Sequences are converted to upper case RNA before folding. Two different quantities are computed from this model, and they treat the pseudoknot differently.

*a. Structure-based potential.* The pairing probabilities $p_{ij}(x,T)$ are those of the unconstrained equilibrium ensemble: for each sequence the partition function is computed over all secondary structures of the Turner model, with no constraint forcing the reference structure, and $\mathcal{S}_{sb}$ then reads off the probabilities of the reference contacts. Because $p_{ij}$ is defined for any pair of positions, all 59 contacts contribute, the six pseudoknot pairs included: the quantity asked of the ensemble is the probability that positions $i$ and $j$ pair, which is well defined even for a pair that the secondary-structure grammar cannot realise jointly with the rest of the reference structure.

*b. Consensus free energy.* The free energy $\Delta G_T(\mathcal{C}_{Azo}|x)$, used only as the control score discussed below, is instead the energy of one prescribed structure, which ViennaRNA evaluates by decomposing it into loops. Since this decomposition does not exist for a pseudoknotted structure, the six pseudoknot pairs are removed and the remaining 53 nested pairs are evaluated; the two positions of each removed pair are treated as unpaired. The contact set used by $\mathcal{S}_{sb}$ is therefore slightly larger than the one entering $\Delta G_T$, which should be kept in mind when the two scores are compared.

#### S3.5. Consensus free energy as an alternative score

Figure S3 shows, for the same set of variants used in Fig. 2 of the main text (317 sequences of the two DCA models of Ref. [1], matched, active at 37°C and 25–30 substitutions from the wild type, of which 165 are active at 60°C), the consensus free energy at 37°C against $\mathcal{H}_{gen}$, with the variants coloured by their activity at 60°C. Active and inactive variants are broadly intermixed along the free-energy axis. Their mean consensus free energies differ only slightly, −52.9 against −49.7 kcal/mol, and the two distributions overlap over essentially their whole range, so no threshold on $\Delta G_T$ separates the two classes in the way a threshold on $\mathcal{S}_{sb}$ does (Fig. 2 of the main text: 86% of the variants with $\mathcal{S}_{sb} < -33$ are active at 60°C, against 34% of those above the threshold). This is the comparison that motivates using the ensemble-based potential rather than the free energy of the target structure: the free energy measures how favourable the consensus fold is in isolation, whereas $\mathcal{S}_{sb}$ measures how much of it survives the competition with alternative folds.

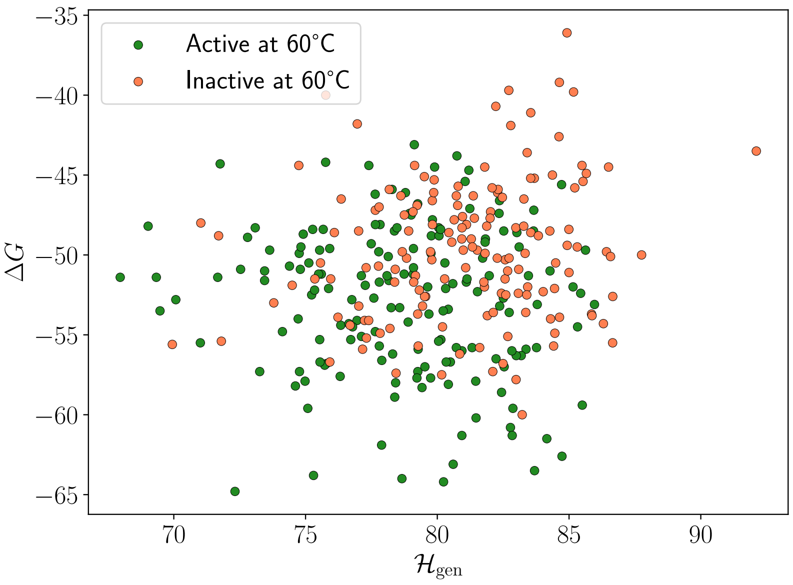


FIG. S3. **Consensus-structure free energy does not separate thermostable variants.** Free energy $\Delta G_{37^\circ C}(\mathcal{C}_{Azo}|x)$ of the consensus structure against the generative score $\mathcal{H}_{gen}$, for the 317 variants of the two DCA models of Ref. [1] that are in the matched subset, active at 37°C and 25–30 substitutions away from the *Azoarcus* wild type. Green: active at 60°C; coral: inactive at 60°C. The two classes overlap along the free-energy axis, in contrast with the separation obtained with $\mathcal{S}_{sb}$ (Fig. 2 of the main text). The pseudoknot pairs are excluded from this free energy, see Sec. S3.4.

#### S3.6. Structure-based potential at 37 and 60°C

All scores used for design and analysis were computed at 37°C. Since the phenotype of interest is self-splicing at 60°C, it is natural to ask whether evaluating the potential at the assay temperature would have changed the picture.

Figure S4 compares the two for the same 317 variants. Raising the folding temperature lowers the probability of every contact, so $\mathcal{S}_{\rm sb}$ shifts upwards by 2.8 units on average (median 2.4), but the two scores rank the variants almost identically: the Pearson correlation is 0.902 and the Spearman correlation 0.883.
The separation between thermostable and non-thermostable variants is likewise unchanged. With the threshold $\mathcal{S}_{\rm sb}(x, 37°{\rm C}) < -33$ used in the main text, 86.1% of the 108 variants below the threshold are active at 60°C against 34.4% of the 209 above it; with the shifted threshold $\mathcal{S}_{\rm sb}(x, 60°{\rm C}) < -30$, the corresponding figures are 86.0% of 100 and 36.4% of 217. Using the potential at the assay temperature would therefore have selected nearly the same sequences; we kept 37°C throughout, which also makes the scores of the designs directly comparable with those of Ref. [1].

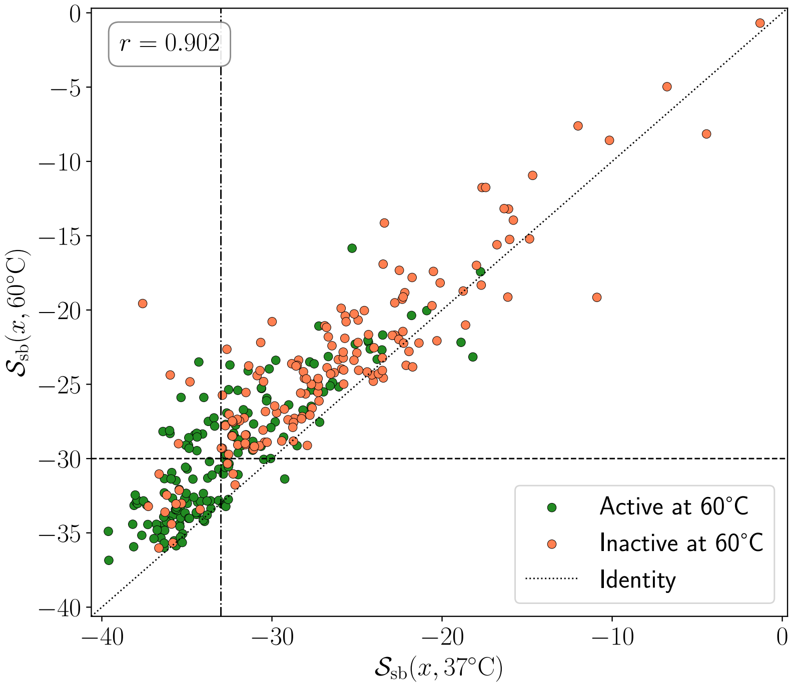


FIG. S4. **The structure-based potential is insensitive to the temperature at which it is evaluated.** $\mathcal{S}_{\rm sb}$ computed at 60°C against $\mathcal{S}_{\rm sb}$ computed at 37°C, for the 317 variants of Fig. S3. Green: active at 60°C; coral: inactive. The dotted line is the identity; points lie above it because contacts are less probable at the higher temperature. The dash-dotted vertical line is the threshold $\mathcal{S}_{\rm sb}(x, 37°{\rm C}) = -33$ used in the main text and the dashed horizontal line its counterpart at 60°C, $\mathcal{S}_{\rm sb} = -30$; the two thresholds select almost the same variants.

### S4. SAMPLING AND LIBRARY CONSTRUCTION

This section gives the proof that the multi-step Monte Carlo scheme of the main text samples the effective Hamiltonian exactly, the parameters and schedule used for each starting sequence and region, and the filters applied to the sampled pools to obtain the synthesised library.

#### S4.1. Detailed balance of the multi-step scheme

Evaluating $\mathcal{S}_{\rm sb}$ requires a partition-function calculation with ViennaRNA and is several orders of magnitude slower than evaluating the Potts energy, so it cannot be recomputed at every attempted substitution. We therefore split the effective Hamiltonian of Eq. (7) of the main text into a “fast” and a “slow” part,

$$\mathcal{H}_{\rm eff}(x \mid x^{\rm st}) = \underbrace{\beta\, \mathcal{H}_{\rm gen}(x) + \lambda_D\, d_H(x^{\rm st}, x)}_{E(x)} + \underbrace{\lambda_S\, \mathcal{S}_{\rm sb}(x)}_{S(x)}, \tag{S7}$$

and alternate many cheap moves that equilibrate $E$ with a single accept/reject test on $S$ applied to the whole block of cheap moves [6]. The target distribution is

$$\pi(x) = \frac{e^{-E(x)-S(x)}}{Z}, \qquad Z = \sum_{x'} e^{-E(x')-S(x')}. \tag{S8}$$

*a. The scheme.* One *slow step* consists of two operations.

(i) Starting from the current state $x^{\mathrm{i}}$, apply a block of fast moves, described by a transition kernel $T(x^{\mathrm{i}} \to x^{\mathrm{f}})$, to obtain a proposal $x^{\mathrm{f}}$. The block uses the fast part of the Hamiltonian only.

(ii) Accept $x^{\mathrm{f}}$ with probability

$$A(x^{\mathrm{i}} \to x^{\mathrm{f}}) = \min\Big(1,\ e^{-\left[S(x^{\mathrm{f}})-S(x^{\mathrm{i}})\right]}\Big), \tag{S9}$$

otherwise return to $x^{\mathrm{i}}$.

*b. The condition on the fast block.* The only property required of the fast block is that it be reversible with respect to the Boltzmann distribution of the fast part alone, $p_E(x) \propto e^{-E(x)}$:

$$e^{-E(x^{\mathrm{i}})}\, T(x^{\mathrm{i}} \to x^{\mathrm{f}}) = e^{-E(x^{\mathrm{f}})}\, T(x^{\mathrm{f}} \to x^{\mathrm{i}}). \tag{S10}$$

Nothing else is assumed about the mechanism inside the block. The fast moves are Gibbs updates: at each elementary move a position $i \in \{1, \dots, L\}$ is drawn uniformly at random and a new nucleotide is emitted from the conditional distribution $\tilde{P}(x_i = a \mid x_{\setminus i}, x^{\mathrm{st}})$ of Eq. (8) of the main text, which is built from the fast parameters $\tilde{h}$ and $\tilde{J}$ only. A single such update leaves $p_E$ invariant and satisfies Eq. (S10); since the position is redrawn at random at every move rather than swept in a fixed order, the elementary kernel is self-adjoint in $L^2(p_E)$, and so is any number of its repetitions. The whole block therefore satisfies Eq. (S10) as well.[1]

*c. Proof.* Detailed balance of the composite move with respect to $\pi$ follows in one line. Using Eq. (S10) and then the elementary identity

$$e^{-S(x^{\mathrm{i}})} \min\Big(1, e^{S(x^{\mathrm{i}})-S(x^{\mathrm{f}})}\Big) = \min\Big(e^{-S(x^{\mathrm{i}})},\ e^{-S(x^{\mathrm{f}})}\Big), \tag{S11}$$

whose right-hand side is manifestly symmetric under $x^{\mathrm{i}} \leftrightarrow x^{\mathrm{f}}$, we obtain

$$\begin{aligned} \pi(x^{\mathrm{i}})\, T(x^{\mathrm{i}} \to x^{\mathrm{f}})\, A(x^{\mathrm{i}} \to x^{\mathrm{f}}) &= \frac{1}{Z}\, e^{-E(x^{\mathrm{i}})} T(x^{\mathrm{i}} \to x^{\mathrm{f}})\, e^{-S(x^{\mathrm{i}})} \min\Big(1, e^{S(x^{\mathrm{i}})-S(x^{\mathrm{f}})}\Big) \\ &= \frac{1}{Z}\, e^{-E(x^{\mathrm{f}})} T(x^{\mathrm{f}} \to x^{\mathrm{i}}) \min\Big(e^{-S(x^{\mathrm{i}})},\ e^{-S(x^{\mathrm{f}})}\Big) \\ &= \pi(x^{\mathrm{f}})\, T(x^{\mathrm{f}} \to x^{\mathrm{i}})\, A(x^{\mathrm{f}} \to x^{\mathrm{i}}). \end{aligned} \tag{S12}$$

The chain is therefore reversible with respect to $\pi$, and since the fast kernel can reach any sequence without gaps in a finite number of moves, it converges to the effective equilibrium distribution (S8). Note that the normalisation $Z$ never enters: only differences of $E$ and of $S$ are evaluated.

*d. Remark: the fast block written out explicitly.* It is instructive to verify Eq. (S12) by writing the fast block as an explicit sequence of moves, which also shows why the intermediate states drop out. Suppose the block consists of $N$ elementary Metropolis moves with symmetric proposals, taking the system along the path $x_0 \to x_1 \to \cdots \to x_N$ with $x_0 = x^{\mathrm{i}}$ and $x_N = x^{\mathrm{f}}$, and write $E_k = E(x_k)$, $S_0 = S(x_0)$, $S_N = S(x_N)$. Detailed balance need only be checked path by path: each forward path is compared with the same path traversed backwards, and summing the resulting equalities over all paths connecting $x^{\mathrm{i}}$ to $x^{\mathrm{f}}$ gives Eq. (S12). The weight of the forward path, times the acceptance of the slow step, is

$$① = \frac{e^{-E_0-S_0}}{Z}\, \chi_{\to}\ \min\big(1, e^{-S_N+S_0}\big), \qquad \chi_{\to} = \prod_{k=1}^{N} \min\big(1, e^{-E_k+E_{k-1}}\big), \tag{S13}$$

[1] The implementation draws one position per move and applies it to all chains at once for efficiency; this correlates the chains with one another but leaves the law of each individual chain, which is the only thing used in the analysis, unchanged.

and that of the reverse path, which visits the same states in the opposite order, is

$$②=\frac{e^{-E_N-S_N}}{Z}\,\chi_{\leftarrow}\;\min\bigl(1,e^{-S_0+S_N}\bigr)\,,\qquad \chi_{\leftarrow}=\prod_{k=1}^{N}\min\bigl(1,e^{E_k-E_{k-1}}\bigr)\,. \tag{S14}$$

Term by term, $\min(1,e^{-a})/\min(1,e^{a})=e^{-a}$ for any real $a$, so the two path weights satisfy

$$\frac{\chi_{\rightarrow}}{\chi_{\leftarrow}}=e^{-E_N+E_0}, \tag{S15}$$

which is Eq. (S10) for this particular path: all intermediate energies $E_1,\ldots,E_{N-1}$ cancel. Substituting Eq. (S15) into Eq. (S13) eliminates $E_0$ in favour of $E_N$ and gives

$$①=\frac{e^{-E_N-S_0}}{Z}\,\chi_{\leftarrow}\;\min\bigl(1,e^{-S_N+S_0}\bigr)\,,\qquad ②=\frac{e^{-E_N-S_N}}{Z}\,\chi_{\leftarrow}\;\min\bigl(1,e^{-S_0+S_N}\bigr)\,. \tag{S16}$$

If $S_0>S_N$ the first minimum equals 1 and the second equals $e^{S_N-S_0}$, so both expressions reduce to $e^{-E_N-S_0}\chi_{\leftarrow}/Z$; if $S_N>S_0$ the roles are exchanged and both reduce to $e^{-E_N-S_N}\chi_{\leftarrow}/Z$. In either case ① = ②, the common value being $e^{-E_N}\min(e^{-S_0},e^{-S_N})\,\chi_{\leftarrow}/Z$, which is Eq. (S11) again.

#### S4.2. Sampling schedule and parameters

For each of the ten starting sequences of Sec. S2 and each of the three target regions, independent chains were initialised at the starting sequence and run for 20 slow steps. Each slow step comprises 10 fast moves, and one fast move is $L=193$ attempted substitutions at randomly chosen modelled positions, so a chain performs 200 fast moves, i.e. 38,600 attempted substitutions, and 20 evaluations of $\mathcal{S}_{\rm sb}$. The local field of the gap symbol was set to a large negative value, so that gaps were never emitted and all designs have the same length as the starting sequence; the three flanking nucleotides 5′-`GUG` and the 3′ `G` are outside the model and were never mutated (Sec. S3.2). $\mathcal{S}_{\rm sb}$ was evaluated at 37°C throughout the sampling (Sec. S3.6).

The weights $\beta$ and $\lambda_S$ were tuned, by trial runs, to place the bulk of the chains in the intended region of the $(\mathcal{H}_{\rm gen},\mathcal{S}_{\rm sb})$ plane; their values are given in Table S2. They depend on the region but not on the starting sequence: the same three parameter sets were used for all ten starts. For the $\uparrow\mathcal{H}_{\rm gen}\downarrow\mathcal{S}_{\rm sb}$ region, $\beta=0$: the generative model is switched off during the sampling and enters only through the selection step below, which is how that region, unfavourable under $\mathcal{H}_{\rm gen}$, is reached at all. Note the sign convention of Eq. (S7): a positive $\lambda_S$ favours low (more negative) $\mathcal{S}_{\rm sb}$, i.e. more consensus pairing, and a negative $\lambda_S$ favours high $\mathcal{S}_{\rm sb}$.

The distance weight $\lambda_D$ was not fixed globally. Because the fast part of the Hamiltonian contains a uniform extra field $+\lambda_D$ on the nucleotide carried by the starting sequence at each position, $\lambda_D$ acts as a single knob controlling the mean distance: the larger it is, the closer the chains stay to $x^{\rm st}$. Its value was therefore tuned case by case, for each starting sequence and region separately, so that the bulk of the histogram of Hamming distances from the starting sequence fell between 5 and 10 substitutions, which is the window later imposed by the filter (Sec. S4.4). The effective value needed to meet that constraint depends on the starting sequence, which is why $\lambda_D$ is not listed in Table S2.

| Region | $\beta$ | $\lambda_S$ | Chains | Slow × fast | Designs/start |
|---|---|---|---|---|---|
| $\downarrow\mathcal{H}_{\rm gen}\downarrow\mathcal{S}_{\rm sb}$ (jointly favoured) | 1.5 | +2.0 | 30,000 | 20 × 10 | 150 |
| $\downarrow\mathcal{H}_{\rm gen}\uparrow\mathcal{S}_{\rm sb}$ (DCA-favoured) | 1.25 | −3.0 | 30,000 | 20 × 10 | 50 |
| $\uparrow\mathcal{H}_{\rm gen}\downarrow\mathcal{S}_{\rm sb}$ (SB-favoured) | 0 | +3.5 | 30,000 | 20 × 10 | 50 |

TABLE S2. **Sampling parameters and schedule.** Score weights of the effective Hamiltonian $\mathcal{H}_{\rm eff}=\beta\mathcal{H}_{\rm gen}+\lambda_S\mathcal{S}_{\rm sb}+\lambda_D d_H(x^{\rm st},\cdot)$ and sampling schedule for each of the three target regions. The values of $\beta$ and $\lambda_S$ were the same for all ten starting sequences; the distance weight $\lambda_D$ was tuned separately for each starting sequence and region and is therefore not listed. "Slow × fast" is the number of slow steps times the number of fast moves per slow step; one fast move is $L=193$ attempted substitutions. "Designs/start" is the number of sequences retained per starting sequence after the filtering of Sec. S4.4, giving $1500+500+500=2500$ designed sequences in the pool (Sec. S5.1).

#### S4.3. Distance constraints

Two distance constraints act on the designs, and they play different roles.
The distance from the starting sequence, $d_H(x^{\rm st}, x)$, is controlled during the sampling by $\lambda_D$ and enforced afterwards by the filter: all retained designs carry between 5 and 10 substitutions with respect to their starting sequence. This window keeps the designs close enough to a sequence of known activity at 37°C that the measured effect can be attributed to the substitutions, while leaving enough room for the two scores to change appreciably.
The distance from the *Azoarcus* wild type, $d_H(x^{\rm wt}, x)$, is not part of the Hamiltonian and is imposed only at the selection stage. Its purpose is to exclude the trivial outcome in which a design is more thermostable than its starting sequence merely because its substitutions revert towards the wild type. The ten starting sequences themselves lie 25–30 substitutions from *Azoarcus* (Table S1), and the nominal budget for the designs is the same 25–30 window.

#### S4.4. Filtering and selection of the library

The pools produced by the chains were filtered, separately for each starting sequence and region, with the criteria of Table I of the main text, namely

- $5 \le d_H(x^{\rm st}, x) \le 10$ and $25 \le d_H(x^{\rm wt}, x) \le 30$ for all regions, the upper limit being raised to 35 for some starting sequences (see below);
- $\mathcal{H}_{\rm gen}(x) < \mathcal{H}_{\rm gen}(x^{\rm st})$ for the two low-$\mathcal{H}_{\rm gen}$ regions, and $\mathcal{H}_{\rm gen}(x) \ge 110$ for the SB-favoured region;
- $\mathcal{S}_{\rm sb}(x, 37°{\rm C}) \le -34$ for the two low-$\mathcal{S}_{\rm sb}$ regions and $\mathcal{S}_{\rm sb}(x, 37°{\rm C}) \ge -10$ for the DCA-favoured region.

The value −33 identified in Fig. 2 of the main text is the threshold that best separates thermostable from non-thermostable variants in the data of Ref. [1]; for the generation we used −34 instead, to keep a margin. Duplicate sequences were removed, and 150 (jointly favoured region) or 50 (the two single-score regions) of the survivors were then drawn uniformly at random, giving the 250 designs per starting sequence that make up the 2500 designed sequences of the pool. The remaining 1000 noise controls and 355 overlap-group sequences were not produced by this procedure; their origin is described in Sec. S5.1.
The $\uparrow \mathcal{H}_{\rm gen} \downarrow \mathcal{S}_{\rm sb}$ region is by far the hardest to populate: the two scores are pushed in opposite directions, and with $\beta = 0$ the chains are driven towards high $\mathcal{H}_{\rm gen}$ rather than towards low $\mathcal{S}_{\rm sb}$ alone. For the starting sequences for which too few candidates passed the $\mathcal{S}_{\rm sb}$ cut, the cut was therefore dropped and the 50 candidates with the lowest $\mathcal{S}_{\rm sb}$ were taken instead. This affects three starting sequences markedly, 4, 7 and 9, whose designs reach $\mathcal{S}_{\rm sb}$ values of −26.9, −27.6 and −24.3, and two more marginally, 1 and 5; over the whole region, 169 of the 500 designs lie above −34. The jointly favoured region, where candidates were plentiful, respects the cut exactly.
The upper end of the distance window from *Azoarcus* was deliberately raised from 30 to 35 for part of the library. For a starting sequence that already lies 30 substitutions from the wild type, staying within 30 while adding 5 to 10 substitutions requires several of them to revert to the wild-type nucleotide, which is precisely what the constraint is meant to prevent; keeping the chains closer than that proved impractical, so the window was widened rather than filled with revertants. This applies to 240 of the 1000 single-score designs, all belonging to starting sequences that are themselves 30 substitutions from *Azoarcus*: the 50 SB-favoured designs of each of starts 6, 7, 8 and 10 lie at 32–35 substitutions, and the 50 DCA-favoured designs of start 6 span 29–35, 40 of them beyond 30. No design in the library exceeds 35, and all 1500 jointly favoured designs respect the original 25–30 window.

### S5. EXPERIMENTAL PROCEDURES

This section reports the full experimental protocol of the pooled self-splicing-like assay, of which the Methods of the main text give a condensed version. The assay follows the design validated in Refs. [1, 2] on RNA pools of up to 18,000 unique molecules. All oligonucleotides are listed in Table S3.

#### S5.1. Library composition

A single pool of 3855 distinct 197-nt sequences was selected for experimental testing. It contains 2500 designed sequences, 250 for each of the ten starting sequences of Sec. S2, split between the three sampled regions of the $(\mathcal{H}_{\rm gen}, \mathcal{S}_{\rm sb})$ plane (Table I of the main text): 150 from the low-$\mathcal{H}_{\rm gen}$/low-$\mathcal{S}_{\rm sb}$ region and 50 from each of the two single-term regions, giving 1500, 500 and 500 sequences in total. In addition, the pool contains 1000 noise-control sequences, more than 100 substitutions away from the *Azoarcus* wild type, which set the background activity level, and the 355 overlap-group sequences of Ref. [1] used as the normalisation reference (Sec. S6).

#### S5.2. RNA preparation

The pool of single-stranded DNA templates corresponding to the designed ribozymes was ordered from Twist Bioscience as part of a pool containing more than 12,000 sequences. Every template is 229 nt long and has the structure 5′–T7 promoter (`TAATACGACTCACTATA`)–tested sequence (197 nt)–exon A (`AATCCGTTGGTGCTG`)–3′; no sequence is therefore added during amplification, and the primers only amplify the construct as synthesised. Both primers span a junction of this construct: F_PCR reproduces the promoter followed by the first three nucleotides `GTG` of the tested sequence, so that the promoter is completed by the template itself and transcription initiates at the 5′-`GUG` of the 197-nt sequence, while R_PCR is the reverse complement of exon A preceded by the last two nucleotides of the tested sequence. Primers and RNA substrates were ordered from Integrated DNA Technologies. Production started from a PCR (18 cycles) with KAPA HiFi HotStart ReadyMix (Roche) and the primers F_PCR and R_PCR (Table S3), followed by clean-up with the NucleoSpin Gel and PCR Cleanup kit (Macherey-Nagel, MN). The RNA pool was transcribed with the HiScribe T7 High Yield RNA Synthesis Kit (New England Biolabs, NEB) for 4 h at 37 °C in a dry bath (MyBlock mini). The template was then removed by treatment with 10 U of DNase I in 1× DNase I buffer (NEB) for 10 min at 37 °C. RNA was recovered by phenol–chloroform extraction followed by precipitation in sodium acetate and ethanol, purified on an 8% urea-PAGE gel, eluted in 0.3 M sodium acetate and precipitated again in sodium acetate/ethanol.

#### S5.3. Self-splicing-like reaction and control

Reactions contained 2 $\mu$M of the RNA pool and 25 $\mu$M each of the substrates S1 and B-S2 in reaction buffer (30 mM EPPS pH 7.5, 60 mM $MgCl_2$), in a final volume of 50 $\mu$L, and were incubated for 1 h at 37 °C or 60 °C. The reaction was stopped with 1.5 $\mu$L of 0.5 M EDTA and purified with the Monarch RNA Cleanup kit (NEB) using adjusted volumes of 100% ethanol (75 $\mu$L) and binding buffer (75 $\mu$L); samples were eluted in 14 $\mu$L of water.
The control sample contained only the 2 $\mu$M RNA pool, with no substrates. It was stopped immediately (0 h) and subjected to the same purification procedure as the reacted sample. Its read frequencies define $f_{\mathrm{ref}}$ and therefore absorb the production biases of the pool (Sec. S6).

#### S5.4. Library preparation and sequencing

Sample preparation for sequencing began with first-strand synthesis using the NEBNext Ultra II RNA First Strand Synthesis Module (NEB). The primer RT_S2, complementary to the substrate S2, was used for the reacted sample, so that only molecules carrying the attached substrate are reverse transcribed, whereas the primer R_PCR, complementary to exon A, was used for the control sample. Second-strand synthesis was carried out with the NEBNext Ultra II Non-Directional RNA Second Strand Synthesis Module (NEB). Samples were cleaned with a 1.2× ratio of NucleoMag NGS Clean-up and Size Select beads (MN) and the final double-stranded DNA concentration was measured with the Qubit dsDNA HS Assay Kit.
Libraries for nanopore sequencing were prepared with the Native Barcoding Kit 24 V14 (SQK-NBD114.24, Oxford Nanopore Technologies, ONT), with AMPure XP bead ratios adjusted to the fragment size; each sample, reacted or control, carried its own barcode and was therefore identified individually in the sequencing run. Libraries were sequenced on PromethION Flow Cells (ONT) in a PromethION 2 Solo device. Basecalling used Dorado (version 1.3.0) with the super-accurate model (ONT). The assignment of reads to designed sequences, the identification of reacted molecules and the normalisation of the resulting frequencies are described in Sec. S6.

### S6. ACTIVITY PROCESSING AND CONTROLS

This section describes how the sequencing reads of the assay of Sec. S5 are turned into the activity values used throughout the paper: how reads are assigned to designed sequences, how the resulting frequencies are normalised, how sequences without a measurable signal are treated, and which controls justify the hit threshold.

TABLE S3. Oligonucleotides used in the assay.

| Name | Sequence | Role |
|---|---|---|
| F_PCR | `TAATACGACTCACTATAGTG` | forward PCR primer containing the T7 promoter |
| R_PCR | `CAGCACCAACGGATTCC` | reverse PCR primer; RT primer of the control sample |
| RT_S2 | `TGAGTTCCGAAGATATTTGAAGTTCC` | RT primer of the reacted sample |
| S1 | `rCrGrCrGrArArUrUrArArCrGrCrGrArCrArArCrArU` | first RNA substrate of the self-splicing-like reaction |
| B-S2 | `rGrGrCrArUrArArCrUrUrCrArArArUrArUrCrUrUrCrGrGrArArCrUrCrA` | second RNA substrate of the self-splicing-like reaction B corresponds to 5′-rGrGrCrArU; the rest to S2 |

#### S6.1. Basecalling and read assignment

Basecalling was performed with Dorado (version 1.3.0) using the super-accurate model (ONT). Each basecalled read is assigned to one of the 3855 synthesised sequences by a segment-matching procedure rather than by alignment: the read is sliced into overlapping segments of 140 nucleotides, each segment is compared with every synthesised sequence, and the read is assigned to the sequence sharing the largest number of segments with it, provided that at least 20 segments are shared. Reads that do not reach this threshold for any sequence are discarded. The procedure is applied independently to the control and to the reacted sample of each temperature.

In the control sample the assigned reads give the frequency $f_{\mathrm{ref}}$ of each sequence in the starting pool, which absorbs the production biases of the synthesis and of the transcription. In the reacted sample only the reads that have lost their 3′ exon A and carry the substrate S2 are retained, so that the assigned reads give the frequency $f_{\mathrm{sel}}$ of molecules that completed both transfer reactions. The raw activity of a sequence at temperature $T$ is

$$b_T(x) = \log_{10} \frac{f_{\mathrm{sel}}(x)}{f_{\mathrm{ref}}(x)}, \tag{S17}$$

with frequencies normalised by the total number of assigned reads of the corresponding sample.

#### S6.2. Read counts and coverage

Table S4 reports, for each group of the pool and for each temperature, the number of assigned reads, the coverage per sequence and the number of sequences without a measurable activity. The control samples yielded $1.14 \times 10^7$ assigned reads at 37 °C and $1.36 \times 10^7$ at 60 °C, that is a median coverage of 2736 and 3233 reads per sequence; the reacted samples yielded $2.7 \times 10^6$ and $3.5 \times 10^6$ assigned reads. Coverage is uniform enough across the three design groups and the overlap group (interquartile ranges within a factor of about three of the median) that no sequence has to be discarded for insufficient reference reads. The noise group is the exception: its median coverage is about half of that of the designs, with a long lower tail, which reflects its lower representation in the synthesised pool rather than the assay itself.

#### S6.3. Missing and undetected activities

Two different situations produce a sequence without a finite activity, and they are treated differently.

A sequence with $f_{\mathrm{ref}} = 0$ has not been observed in the control sample at all. Its activity is *missing*: no value is assigned and the sequence is excluded from every statistic at that temperature. In this experiment this happens for a single sequence, one noise control at 60 °C, so the only quantities affected by missing data are the noise-control fractions at 60 °C, which refer to the 999 noise controls with control reads.

A sequence with $f_{\mathrm{ref}} > 0$ but $f_{\mathrm{sel}} = 0$ has been observed in the pool but never among the reacted molecules. Its activity is *undetected*: Eq. (S17) gives $a_T = -\infty$, and the sequence is classified as inactive. Undetected sequences are kept in all counts and hit rates with their value treated as $-\infty$; the isolated leftmost bin used for them in Figs. 4 and 5 of the main text is a display convention and is not a quantitative activity value.

TABLE S4. **Sequencing counts, coverage and sequences without measured activity.** For each group of the pool (Sec. S5.1) and each temperature: $N$, the number of distinct sequences; the assigned reads of the control sample ($f_{\mathrm{ref}}$) and of the reacted sample ($f_{\mathrm{sel}}$), in units of $10^3$ reads; the median number of control reads per sequence, with the interquartile range; "missing", the number of sequences with no control read, for which no activity is defined; and "undetected", the number of sequences with control reads but no reacted read, whose activity is a lower bound. Read counts as released with the paper, rounded to the nearest integer.

| Group | $T$ (°C) | $N$ | Control reads | Reacted reads | Reads/seq. (IQR) | Missing | Undetected |
|---|---|---|---|---|---|---|---|
| $\downarrow \mathcal{H}_{\mathrm{gen}} \downarrow \mathcal{S}_{\mathrm{sb}}$ | 37 | 1500 | 4762 | 2206 | 3126 (2304–3909) | 0 | 4 |
| | 60 | | 5624 | 2260 | 3688 (2662–4666) | 0 | 5 |
| $\downarrow \mathcal{H}_{\mathrm{gen}} \uparrow \mathcal{S}_{\mathrm{sb}}$ | 37 | 500 | 1799 | 112 | 3487 (2362–4547) | 0 | 18 |
| | 60 | | 2178 | 45 | 4206 (2836–5578) | 0 | 34 |
| $\uparrow \mathcal{H}_{\mathrm{gen}} \downarrow \mathcal{S}_{\mathrm{sb}}$ | 37 | 500 | 1773 | 6 | 3246 (2250–4302) | 0 | 91 |
| | 60 | | 2147 | 3 | 3934 (2772–5356) | 0 | 149 |
| Noise control | 37 | 1000 | 2385 | 2 | 1182 (410–3207) | 0 | 710 |
| | 60 | | 2906 | 1 | 1434 (436–3905) | 1 | 743 |
| Reference (overlap) | 37 | 355 | 661 | 405 | 1865 (1288–2417) | 0 | 6 |
| | 60 | | 766 | 1206 | 2159 (1510–2761) | 0 | 7 |
| All | 37 | 3855 | 11 381 | 2731 | 2736 (1402–3866) | 0 | 829 |
| | 60 | | 13 620 | 3516 | 3233 (1663–4657) | 1 | 938 |

Treating them as inactive rather than discarding them matters, because they are not rare and they are concentrated in the groups with little activity: 0.3% of the jointly optimised designs are undetected at 37 °C against 18.2% of the structure-only designs and 71.0% of the noise controls, and the corresponding figures at 60 °C are 0.3%, 29.8% and 74.4% (Table S4). Discarding them would therefore remove most of the inactive sequences and inflate the apparent activity of the weakest groups.

#### S6.4. Reference group and normalisation

Raw activities are not comparable between samples, because $f_{\mathrm{sel}}$ and $f_{\mathrm{ref}}$ are frequencies within libraries of different sizes and different reaction yields. They are placed on a common scale with the reference group: the 355 sequences of Ref. [1] included in every pool (Sec. S5.1). The normalised activity is

$$a_T(x) = b_T(x) - \bar{b}_{\mathrm{ref},T}, \qquad \bar{b}_{\mathrm{ref},T} = \frac{1}{|R_T|} \sum_{x' \in R_T} b_T(x'), \tag{S18}$$

where $R_T$ is the set of reference sequences with a detected activity at temperature $T$ (349 of 355 at 37 °C and 348 at 60 °C; the remaining ones are undetected and are excluded from the mean). The subtracted means are $\bar{b}_{\mathrm{ref},37} = 0.031$ and $\bar{b}_{\mathrm{ref},60} = -0.187$. By construction the detected reference sequences have mean normalised activity zero at both temperatures, so $a_T$ measures the enrichment of a sequence relative to the geometric mean enrichment of the reference group, in decades.

The reference group was not chosen to be homogeneous: it spans 10 to 30 substitutions from the *Azoarcus* wild type, generative energies from about 70 to 165 and structure-based potentials from −49 to −1 (Fig. S7a–c), while being composed of sequences that were active in the experiment of Ref. [1]. It therefore provides a stable and reproducible scale rather than a single reference point, which is what is needed to compare pools assayed in separate experiments. Its normalised activity distributions are shown in Fig. S7d,e: at 37 °C they are concentrated within about half a decade of the reference mean, and at 60 °C they spread out, as expected for sequences that were selected for activity at 37 °C only.

A sequence is called a *hit* when $a_T \geq -1$, that is when its enrichment is at least 10% of the geometric mean enrichment of the reference group. This is a relative criterion, deliberately not a $p$-value against the noise background; the next subsection shows why.

#### S6.5. Noise controls and the hit threshold

The natural way to define an activity threshold would be to model the measurements of sequences that cannot splice and to cut at a chosen tail probability. The 1000 noise controls, at 100 to 149 substitutions from the wild type, are the relevant background: no functional sequence was found at such distances in Ref. [1]. Their raw activity distributions, Fig. S5a,b, are however clearly not Gaussian. Both are right-skewed (skewness 0.67 at 37 °C and 0.58 at 60 °C) with an extended upper tail of spuriously high apparent activity, and a Shapiro–Wilk test rejects normality at both temperatures ($p = 2 \times 10^{-5}$ and $p = 3 \times 10^{-4}$). A threshold taken from the 1% tail of a Gaussian fitted to these measurements would fall at $b = -1.50$ and $b = -2.19$ respectively, in a region where the empirical distribution and the Gaussian disagree by a large factor. We therefore did not derive the threshold from a background model.
The noise group is instead used the other way round, to check the threshold defined in Sec. S6.4. Of the 1000 noise controls, 290 at 37 °C and 256 at 60 °C have a detected activity, the rest being undetected (one of them has no control read at 60 °C and is missing rather than undetected). Their normalised activities, Fig. S5c,d, lie entirely below the hit threshold: the largest value is −1.00 at 37 °C and −1.65 at 60 °C, and the 99.9th percentile is −1.05 and −1.66. Not one noise control is a hit at either temperature. The threshold $a_T \geq -1$ is therefore conservative with respect to the measured background, and the activity of a hit is very unlikely to arise from measurement noise alone. At 37 °C the margin is thin, a single noise sequence sitting 0.002 decades below the threshold, so the threshold should be read as the edge of the noise distribution rather than as a comfortable separation from it.

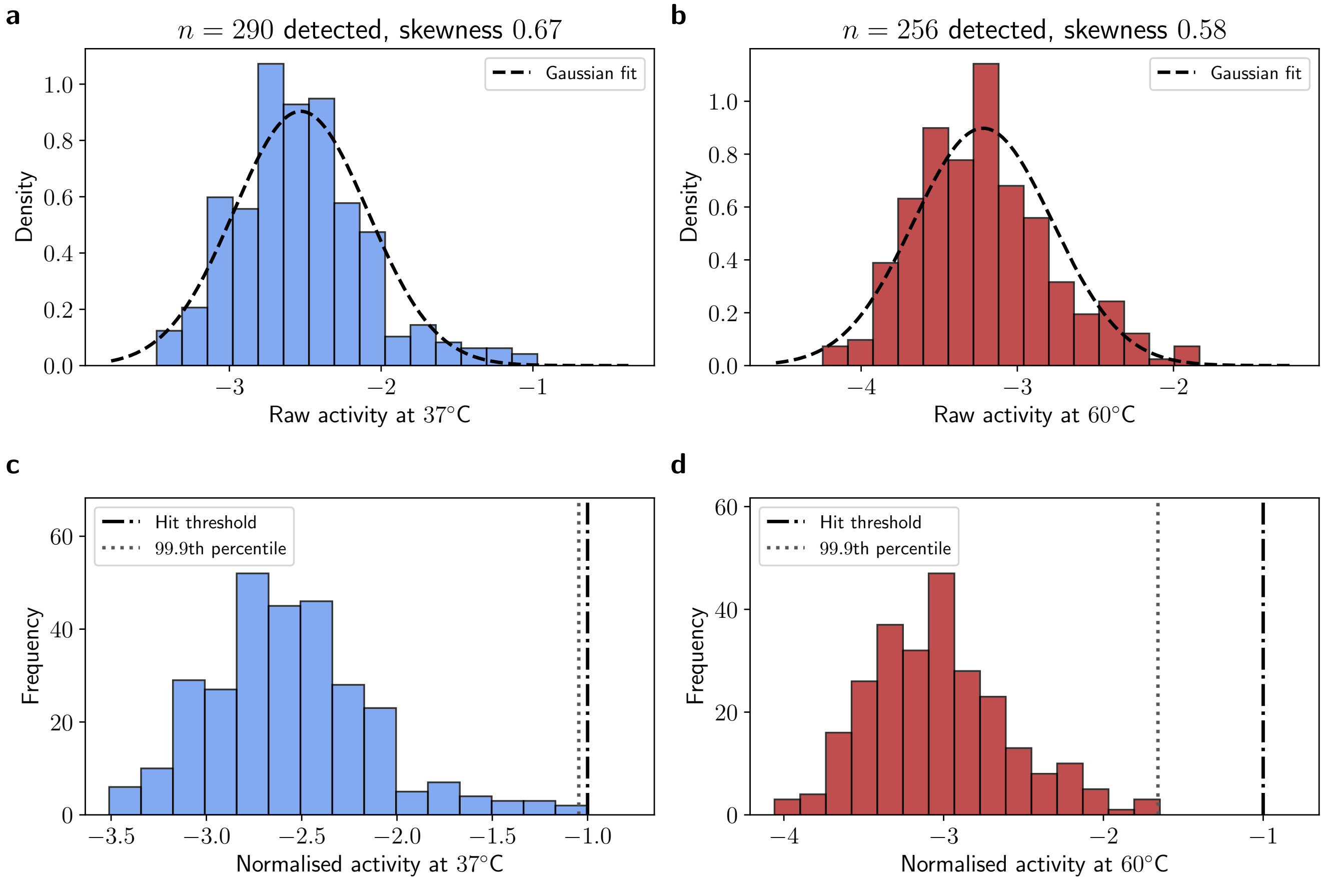


FIG. S5. **Noise controls and the hit threshold.** **a**,**b**: raw activity $b_T$ of the noise-control sequences with a detected activity, at 37 °C ($n = 290$) and at 60 °C ($n = 256$), with the Gaussian of the same mean and variance (dashed). Both distributions are right-skewed and are not described by the Gaussian background model that would be needed for a $p$-value threshold. **c**,**d**: the same measurements after reference normalisation, Eq. (S18). The dash-dotted line is the hit threshold $a_T = -1$ and the dotted line the 99.9th percentile of the noise distribution. No noise control reaches the threshold at either temperature. Sequences with an undetected activity (710 and 743 of the 1000 and 999 noise controls with control reads at the two temperatures ) are not shown.

#### S6.6. The activity cutoffs of the previous dataset

The same reasoning justifies the two cutoffs that we take from Ref. [1] whenever a variant of that dataset is called active: $a_{37} > -2.76$ and $a_{60} > -2/\ln 10 \simeq -0.87$ (Secs. S2.1 and S2.3). They are presented as illustrative in Ref. [1], but they are not arbitrary, because that dataset carries its own background: variants far enough from the *Azoarcus* wild type cannot fold into a functional ribozyme, so whatever activity is still measured for them is the background of the assay. Figure S6 shows that both cutoffs sit at the top edge of that background.

At 37 °C (Fig. S6a) the 99th percentile of the measured activity falls steadily with the distance from the wild type and then stops falling, flattening onto the cutoff beyond about 60 substitutions: of the 3396 variants with a detected activity past that distance, only 0.8% exceed −2.76, which is their 99.2nd percentile. The fraction of variants above the cutoff drops from 76% within 20 substitutions to 45%, 26%, 2.4% and 0.2% in the following bins of 20. The distribution of the whole dataset (Fig. S6c) tells the same story from the other side: it is clearly bimodal, with an inactive mode near −4 and an active mode near −0.2, and the cutoff lies in the valley between them.

At 60 °C (Fig. S6b,d) the same pattern holds, with the per-bin 99th percentile reaching the cutoff at about the same distance, and the fraction above it dropping from 85% to 59%, 22%, 4.9% and 0% across the same bins. The evidence is thinner here: the 60 °C condition was applied mostly to variants close to the wild type, so only 44 variants with a detected activity lie beyond 60 substitutions, of which 4.5% exceed the cutoff. The value itself is a round number in the natural-logarithm units of Ref. [1], −2, rather than a quantity estimated from the data.

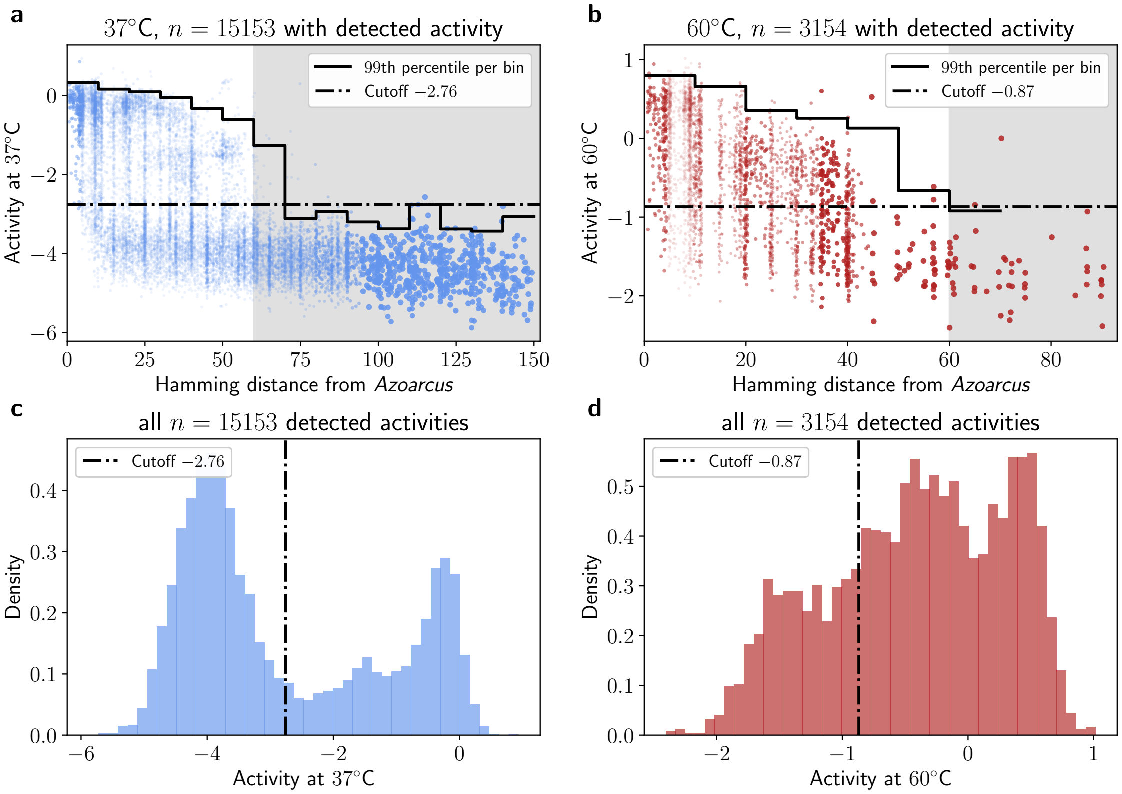


FIG. S6. **The activity cutoffs of Ref. [1] against the distance from the wild type. a,b**: measured activity of every variant of Ref. [1] with a detected activity, at 37 °C and at 60 °C, against its Hamming distance from the *Azoarcus* wild type (points jittered horizontally). The dash-dotted line is the cutoff and the step curve the 99th percentile of the activity in bins of 10 substitutions, computed where the bin holds at least 20 variants. The shaded region marks the variants more than 60 substitutions away, which are too distant to splice and whose measurements therefore define the background of the assay; the step curve flattens onto the cutoff there. **c,d**: the distribution of the same activities, with the cutoff. At 37 °C it falls in the valley between the inactive and the active mode of a clearly bimodal distribution; at 60 °C, where the distribution has no comparable valley, it sits on the shoulder of the active population.

#### S6.7. Agreement with the previous experiment

The reference group also gives an external check of the whole pipeline, because its sequences were measured again here after having been measured in Ref. [1]. Of the 355 reference sequences, 144 are present in the published dataset of Ref. [1] and 138 of them have a detected activity at 37 °C in both experiments. Their activities agree well (Fig. S7f): the Pearson correlation is 0.96 and the Spearman correlation 0.76, the lower rank correlation reflecting the narrow range spanned by the bulk of the group rather than a disagreement. Two independent assays, performed on different pools and analysed with different pipelines, therefore place the same sequences on compatible activity scales, which supports using the reference group as the common scale. The comparison cannot be repeated at 60 °C, where the sequences of this group have no measurement in Ref. [1].

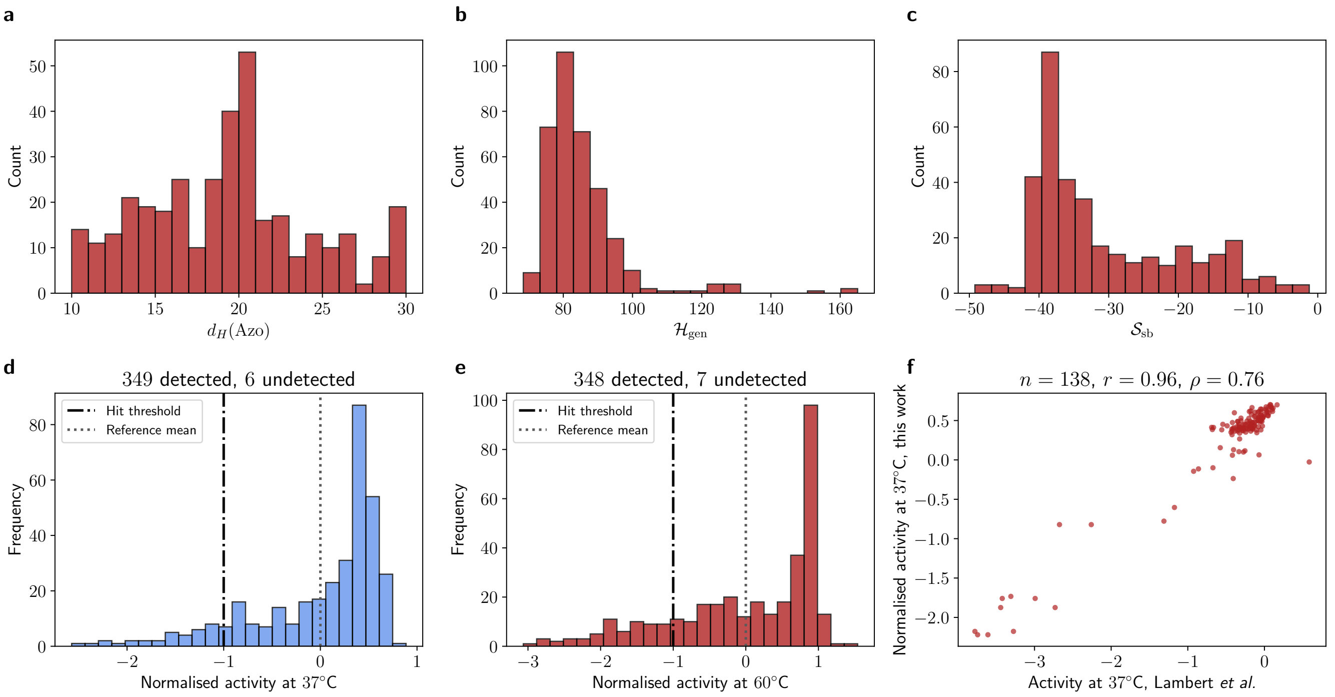


FIG. S7. **The 355-sequence reference group.** **a**–**c**: coverage of the reference group in Hamming distance from the *Azoarcus* wild type, in generative energy $\mathcal{H}_{\rm gen}$ and in structure-based potential $\mathcal{S}_{\rm sb}$ evaluated at 37 °C. **d**,**e**: reference-normalised activities of the group at the two temperatures; the dotted line is the reference mean, zero by construction, and the dash-dotted line the hit threshold $a_T = -1$. Undetected sequences (6 and 7) are not shown. **f**: activity at 37 °C measured here against the activity measured in Ref. [1], for the 138 reference sequences with a detected activity in both experiments. The two scales differ by the normalisation of Eq. (S18), which is why the points do not lie on the diagonal.

### S7. RESULTS BY STARTING SEQUENCE

The main text reports the outcome of the assay pooled over the ten starting sequences. This section reports the same quantities separately for each starting sequence and each sampling region, so that the pooled numbers can be checked against the variability between the ten independent design problems. Table S5 gives the numbers and Fig. S8 shows the underlying distributions.

#### S7.1. Quantities reported

Every starting sequence contributes 150 designs to the jointly favoured region ($\downarrow \mathcal{H}_{\rm gen} \downarrow \mathcal{S}_{\rm sb}$) and 50 to each of the two single-score regions, for the 2500 designs of the library (Sec. S4.4). Activities are the reference-normalised $a_T$ of Eq. (S18) and a hit has $a_T \geq -1$, as in Sec. S6.5. Undetected activities are kept at $-\infty$ in every count, quantile and hit rate, as in Sec. S6.3. They are also listed separately in Table S5, because they are the dominant contribution to the low activity levels of the structure-only region. Two further quantities summarise the temperature response: the *retention*, the fraction of the designs that are hits at 37 °C and are hits again at 60 °C, and the median of $\Delta = a_{60} - a_{37}$ over the same subset of designs, that is the activity change used in Fig. 5 of the main text restricted to one starting sequence and one region.

The ten starting sequences themselves were not re-synthesised in this library (Sec. S2.2), so they have no row of their own here; their baseline activities, measured in Ref. [1], are those of Table S1. All ten are active at 37 °C and have no detected activity at 60 °C, which is the baseline against which the numbers below should be read. Table S5 is also supplied as a machine-readable file, alongside the per-sequence activities and counts.

#### S7.2. The ordering of the three regions holds for every starting sequence

The ordering reported for the pooled library is reproduced by each of the ten starting sequences separately: at both temperatures the jointly favoured region has the highest hit rate, the DCA-favoured region ($\downarrow \mathcal{H}_{\rm gen} \uparrow \mathcal{S}_{\rm sb}$) an intermediate one and the structure-only region ($\uparrow \mathcal{H}_{\rm gen} \downarrow \mathcal{S}_{\rm sb}$) an almost vanishing one (Fig. S8a,b). At 60 °C the jointly favoured region exceeds the DCA-favoured region for 10 starting sequences out of 10 (two-sided Wilcoxon signed-rank test on the ten paired hit rates, $p = 2 \times 10^{-3}$), with hit rates of 40.7%–99.3% against 2.0%–20.0%. The pooled comparison of the main text is therefore not driven by a subset of the starting sequences.

The same holds for the temperature response. The median $\Delta$ of the jointly favoured region exceeds that of the DCA-favoured region for all ten starting sequences, by 0.15 to 1.02 decades (pooled value: 0.6 decades, Fig. 5 of the main text), and the difference between the two distributions of $\Delta$ is significant for nine of the ten starting sequences (one-sided Mann–Whitney test, $p < 0.02$; the exception is start 4, $p = 0.13$, where the DCA-favoured region has only 14 hits at 37 °C to compare). Retention follows the same ordering: 53%–99% of the jointly favoured designs that are hits at 37 °C are still hits at 60 °C, against 6%–64% in the DCA-favoured region.

#### S7.3. Variability between starting sequences

The absolute level of the response varies considerably between starting sequences. In the jointly favoured region the hit rate at 37 °C ranges from 65.3% (start 2) to 96.7% (start 8) and at 60 °C from 40.7% (start 2) to 99.3% (start 8); starts 1, 2 and 7 are the three least successful at the high temperature and starts 8, 9 and 4 the three most successful.

#### S7.4. The structure-only region

The structure-only region produces almost no activity for any starting sequence: 6 of its 500 designs are hits at 37 °C and 2 at 60 °C, and they belong to only three starting sequences (1, 4 and 5). These few designs sit at the edge of the region: their generative energies, 110.0 to 116.5, are at the lower boundary $\mathcal{H}_{\rm gen} > 110$ of the region, and four of the six have $\mathcal{S}_{\rm sb}$ between $-30.2$ and $-27.0$, that is above the threshold $-34$ actually used for generation, so they come from the starting sequences for which too few candidates passed that threshold and the best available ones were taken instead (Sec. S4.4). The handful of active designs of this region is therefore found where the generative energy is least unfavourable, and not where the structure-based potential is most favourable.

The undetected activities are also concentrated in this region and are unevenly distributed over the starting sequences: 91 of its 500 designs are undetected at 37 °C and 149 at 60 °C, of which 25 and 37 belong to start 6 alone, whose structure-only designs are undetected in such a proportion at 60 °C (37 of 50) that even their upper quartile falls among the undetected ones. For comparison, the jointly favoured region has 4 and 5 undetected designs in total out of 1500.

TABLE S5. **Results by starting sequence and sampling region.** For each starting sequence and each of the three regions: $N$, the number of designs; "undet.", the number of designs with an undetected activity at 37 and at 60 °C; "top 25%", the reference-normalised activity delimiting the highest-activity quartile at the two temperatures (the 75th percentile, as in Table II of the main text); the hit rate at the two temperatures, in per cent; the retention, the percentage of the designs that are hits at 37 °C which are hits again at 60 °C; the median $\tilde{\Delta}$ of $\Delta = a_{60} - a_{37}$ over those same designs; and the mean changes $\Delta\Delta G$ (in kcal mol$^{-1}$) and Δ H-bonds of the Turner free energy and of the number of hydrogen bonds of the consensus structure, relative to the corresponding starting sequence. Quantiles and hit rates include the undetected designs as $-\infty$. An asterisk marks an upper quartile that is itself $-\infty$: there more than a quarter of the designs of that region are undetected, so the quartile falls among them and the measurement only bounds it from above. A dash marks a quantity that is undefined because the region has no hit at 37 °C for that starting sequence. The last block repeats the three regions pooled over the ten starting sequences, as in Table II of the main text.

| Start | Region | $N$ | Undet. (37/60) | Top 25% (37 °C) | Top 25% (60 °C) | Hit 37 (%) | Hit 60 (%) | Retention (%) | $\tilde{\Delta}$ | $\Delta\Delta G$ | Δ H-bonds |
|---|---|---|---|---|---|---|---|---|---|---|---|
| 1 | $\downarrow\mathcal{H}_{\rm gen}\downarrow\mathcal{S}_{\rm sb}$ | 150 | 0/0 | −0.10 | −0.57 | 95.3 | 50.7 | 53.1 | −0.71 | −5.08 | 9.71 |
| | $\downarrow\mathcal{H}_{\rm gen}\uparrow\mathcal{S}_{\rm sb}$ | 50 | 0/5 | −0.77 | −1.75 | 38.0 | 4.0 | 10.5 | −1.10 | 0.34 | 2.26 |
| | $\uparrow\mathcal{H}_{\rm gen}\downarrow\mathcal{S}_{\rm sb}$ | 50 | 11/13 | −2.61 | −2.55 | 2.0 | 0.0 | 0.0 | −1.33 | 1.36 | 0.12 |
| 2 | $\downarrow\mathcal{H}_{\rm gen}\downarrow\mathcal{S}_{\rm sb}$ | 150 | 2/3 | −0.13 | −0.71 | 65.3 | 40.7 | 56.1 | −0.68 | −4.76 | 6.66 |
| | $\downarrow\mathcal{H}_{\rm gen}\uparrow\mathcal{S}_{\rm sb}$ | 50 | 2/3 | −0.86 | −1.92 | 28.0 | 10.0 | 35.7 | −1.19 | −0.74 | 1.48 |
| | $\uparrow\mathcal{H}_{\rm gen}\downarrow\mathcal{S}_{\rm sb}$ | 50 | 4/7 | −2.11 | −2.60 | 0.0 | 0.0 | — | — | 1.78 | −3.22 |
| 3 | $\downarrow\mathcal{H}_{\rm gen}\downarrow\mathcal{S}_{\rm sb}$ | 150 | 0/1 | 0.26 | 0.54 | 92.7 | 80.0 | 86.3 | 0.09 | −2.44 | 4.51 |
| | $\downarrow\mathcal{H}_{\rm gen}\uparrow\mathcal{S}_{\rm sb}$ | 50 | 4/5 | −0.63 | −1.40 | 40.0 | 6.0 | 15.0 | −0.93 | 4.05 | −2.26 |
| | $\uparrow\mathcal{H}_{\rm gen}\downarrow\mathcal{S}_{\rm sb}$ | 50 | 12/17 | −2.32 | −2.81 | 0.0 | 0.0 | — | — | 2.08 | −1.60 |
| 4 | $\downarrow\mathcal{H}_{\rm gen}\downarrow\mathcal{S}_{\rm sb}$ | 150 | 0/0 | 0.44 | 0.45 | 90.7 | 85.3 | 93.4 | −0.11 | −4.69 | 3.98 |
| | $\downarrow\mathcal{H}_{\rm gen}\uparrow\mathcal{S}_{\rm sb}$ | 50 | 1/4 | −0.90 | −1.19 | 28.0 | 18.0 | 64.3 | −0.25 | −1.60 | −0.04 |
| | $\uparrow\mathcal{H}_{\rm gen}\downarrow\mathcal{S}_{\rm sb}$ | 50 | 3/6 | −1.84 | −2.26 | 6.0 | 4.0 | 33.3 | −2.14 | 4.34 | −4.50 |
| 5 | $\downarrow\mathcal{H}_{\rm gen}\downarrow\mathcal{S}_{\rm sb}$ | 150 | 1/0 | 0.03 | −0.20 | 95.3 | 69.3 | 72.7 | −0.51 | −3.66 | 3.66 |
| | $\downarrow\mathcal{H}_{\rm gen}\uparrow\mathcal{S}_{\rm sb}$ | 50 | 0/0 | −0.83 | −1.62 | 32.0 | 2.0 | 6.2 | −1.15 | 3.45 | −2.10 |
| | $\uparrow\mathcal{H}_{\rm gen}\downarrow\mathcal{S}_{\rm sb}$ | 50 | 5/20 | −2.21 | −2.94 | 2.0 | 0.0 | 0.0 | −2.05 | 1.99 | −2.10 |
| 6 | $\downarrow\mathcal{H}_{\rm gen}\downarrow\mathcal{S}_{\rm sb}$ | 150 | 0/0 | 0.63 | −0.19 | 86.7 | 73.3 | 82.3 | −0.42 | −0.61 | 0.97 |
| | $\downarrow\mathcal{H}_{\rm gen}\uparrow\mathcal{S}_{\rm sb}$ | 50 | 4/9 | −1.16 | −2.07 | 18.0 | 2.0 | 11.1 | −1.14 | 5.14 | −5.18 |
| | $\uparrow\mathcal{H}_{\rm gen}\downarrow\mathcal{S}_{\rm sb}$ | 50 | 25/37 | −2.71 | $-\infty^*$ | 0.0 | 0.0 | — | — | 2.92 | −3.46 |
| 7 | $\downarrow\mathcal{H}_{\rm gen}\downarrow\mathcal{S}_{\rm sb}$ | 150 | 1/1 | 0.00 | −0.47 | 76.0 | 52.0 | 66.7 | −0.41 | −9.95 | 6.62 |
| | $\downarrow\mathcal{H}_{\rm gen}\uparrow\mathcal{S}_{\rm sb}$ | 50 | 0/0 | −1.07 | −1.90 | 20.0 | 2.0 | 10.0 | −1.18 | −2.71 | −0.82 |
| | $\uparrow\mathcal{H}_{\rm gen}\downarrow\mathcal{S}_{\rm sb}$ | 50 | 4/4 | −2.30 | −2.65 | 0.0 | 0.0 | — | — | −0.29 | −3.00 |
| 8 | $\downarrow\mathcal{H}_{\rm gen}\downarrow\mathcal{S}_{\rm sb}$ | 150 | 0/0 | 0.70 | 0.35 | 96.7 | 99.3 | 99.3 | −0.34 | −3.81 | 7.47 |
| | $\downarrow\mathcal{H}_{\rm gen}\uparrow\mathcal{S}_{\rm sb}$ | 50 | 1/0 | −0.21 | −1.25 | 48.0 | 20.0 | 33.3 | −1.28 | 2.69 | 0.66 |
| | $\uparrow\mathcal{H}_{\rm gen}\downarrow\mathcal{S}_{\rm sb}$ | 50 | 8/5 | −1.84 | −2.29 | 0.0 | 0.0 | — | — | 1.42 | −0.68 |
| 9 | $\downarrow\mathcal{H}_{\rm gen}\downarrow\mathcal{S}_{\rm sb}$ | 150 | 0/0 | 0.15 | −0.06 | 96.0 | 88.0 | 89.6 | −0.28 | −4.76 | 8.41 |
| | $\downarrow\mathcal{H}_{\rm gen}\uparrow\mathcal{S}_{\rm sb}$ | 50 | 2/2 | −0.96 | −1.68 | 30.0 | 2.0 | 6.7 | −0.79 | 0.36 | 1.80 |
| | $\uparrow\mathcal{H}_{\rm gen}\downarrow\mathcal{S}_{\rm sb}$ | 50 | 9/21 | −2.01 | −2.47 | 0.0 | 0.0 | — | — | 2.58 | −1.96 |
| 10 | $\downarrow\mathcal{H}_{\rm gen}\downarrow\mathcal{S}_{\rm sb}$ | 150 | 0/0 | 0.20 | 0.04 | 94.7 | 86.0 | 90.1 | −0.22 | −1.54 | 6.07 |
| | $\downarrow\mathcal{H}_{\rm gen}\uparrow\mathcal{S}_{\rm sb}$ | 50 | 4/6 | −0.71 | −1.44 | 40.0 | 12.0 | 30.0 | −0.74 | 3.58 | 0.38 |
| | $\uparrow\mathcal{H}_{\rm gen}\downarrow\mathcal{S}_{\rm sb}$ | 50 | 10/19 | −2.11 | −2.72 | 0.0 | 0.0 | — | — | 2.96 | −1.42 |
| All | $\downarrow\mathcal{H}_{\rm gen}\downarrow\mathcal{S}_{\rm sb}$ | 1500 | 4/5 | 0.25 | 0.02 | 88.9 | 72.5 | 79.9 | −0.37 | −4.13 | 5.81 |
| | $\downarrow\mathcal{H}_{\rm gen}\uparrow\mathcal{S}_{\rm sb}$ | 500 | 18/34 | −0.83 | −1.56 | 32.2 | 7.8 | 23.0 | −0.97 | 1.46 | −0.38 |
| | $\uparrow\mathcal{H}_{\rm gen}\downarrow\mathcal{S}_{\rm sb}$ | 500 | 91/149 | −2.12 | −2.61 | 1.0 | 0.4 | 20.0 | −2.05 | 2.11 | −2.18 |

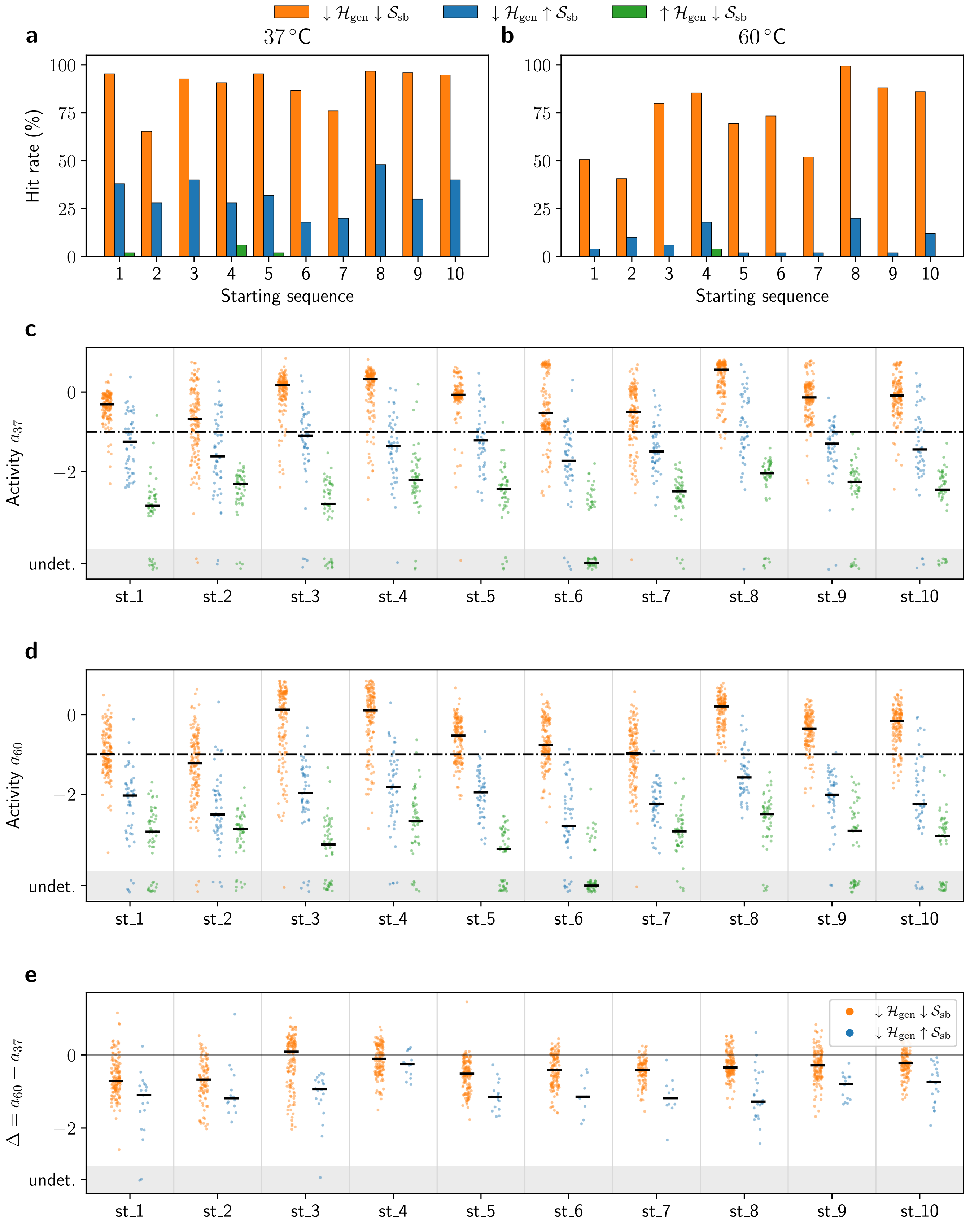


FIG. S8. **Results by starting sequence.** Colours denote the three sampling regions: jointly favoured (orange), DCA-favoured (blue) and structure-only (green). **a**,**b**: hit rate of each region for each starting sequence, at 37 and at 60 °C. **c**,**d**: reference-normalised activities of the individual designs at 37 and at 60 °C, one column per starting sequence; the horizontal bars are the medians, including the undetected designs, and the dash-dotted line is the hit threshold $a_T = -1$. **e**: activity change $\Delta = a_{60} - a_{37}$ of the designs that are hits at 37 °C, for the jointly favoured and the DCA-favoured regions; the structure-only region is omitted because it has at most three such designs per starting sequence. Designs with an undetected activity are drawn in the shaded band at the bottom of each panel, which is a display convention and not an activity value.

### S8. MD METHODS AND ROBUSTNESS CHECKS

This section describes the molecular dynamics (MD) simulations and the analyses of the wild-type *Azoarcus* group I intron (WT), Start–10, and the two Start–10-derived variants considered in the main text. We refer to the orange variant as the *jointly favoured* variant and to the blue variant as the *DCA favoured* variant throughout. Residue numbers refer to the full 197-nucleotide construct. DCA alignment positions 1–193 correspond to construct positions 4–196.

The MD comparison was motivated by the different responses to heating of two Start–10-derived variants with the same nine-substitution budget and similar DCA energies. Their normalized $\log_{10}$ activities change from −0.61 to 0.41 (jointly favored) and from −0.90 to −2.43 (DCA-favored) between 37 and 60 °C. The systems remain broadly compact during the analyzed 0–300 ns interval, so the comparison concerns differences in motion and local geometry within the sampled folded conformations rather than observed global unfolding. At 60 °C, the jointly favored variant shows lower sequence-wide displacements, including at terminal ΩG; the quantitative displacement analysis and time-window checks are already reported in Sec. S8.4 and Fig. S9. This motivated the distance-based analysis of the catalytic pocket, whose definitions and reference geometry are given in Secs. S8.5 and S8.6.

#### S8.1. Sequences and structure prediction

The jointly favoured variant is candidate `start10_lowDCA_lowSB_031`; the DCA favoured variant is candidate `start10_lowDCA_highSB_007`. Each differs from Start–10 by nine substitutions (Table S6). Their generative energies are $\mathcal{H}_{\text{gen}} = 76.7$ and 73.9, respectively, and their structure-based potentials, evaluated at 37 °C, are $\mathcal{S}_{\text{sb}} = -36.2$ and −9.6.

Sequence-specific starting structures were predicted with AlphaFold 3 (version 3.0.1) [7]. Fifteen models were generated per sequence, and the model with the highest ranking score was retained for simulation. The models recovered the same overall architecture, with local differences, including at the 3′ terminus.

#### S8.2. System preparation and simulation conditions

Simulations were performed with GROMACS 2025.1 [8], using the `amber14sb_99bsc0-cOL3_OPC` force-field distribution, which includes the parmbsc0 and χOL3 nucleic-acid corrections [9, 10], and explicit OPC water [11]. Each RNA was placed in a periodic dodecahedral box with a minimum RNA-to-box clearance of 1.2 nm. The nominal salt concentrations were 0.15 M KCl and 5 mM $MgCl_2$.

The two temperatures were 310 and 333 K, corresponding approximately to 37 and 60 °C. Temperature was controlled with the V-rescale thermostat [12]; pressure coupling used the Parrinello–Rahman barostat [13]. Bonds involving hydrogen were constrained with LINCS [14]. The equilibration protocol comprised 0.5 ns in the NVT ensemble, 1 ns in the NPT ensemble, and a further 10 ns of unrestrained equilibration.

- **Software.** GROMACS 2025.1 (local build), GPU offload of non-bonded, PME and bonded terms, 32 OpenMP threads.
- **Force field and water.** AMBER14SB with $\chi_{\text{OL3}}$/bsc0 RNA corrections (`amber14sb_99bsc0-cOL3_OPC`); OPC four-site water model (water-model entry 6 of `watermodels.dat`).
- **System.** RNA of 197 nucleotides, net charge $-196\,e$; rhombic dodecahedral box with a 1.2 nm solute–boundary distance; 57 587 OPC water molecules; 237 255 atoms in total; production box vectors 13.64 × 13.64 × 9.64 nm.
- **Terminal chemistry.** The 5′ terminus was converted to a free hydroxyl by deleting `OP3` and the terminal phosphate atoms (`P`, `OP1`, `OP2`), giving the AMBER `RG5` cap (`O5'–H5T`); the 3′ terminus is a free hydroxyl (`RG3` cap, `O3'–H3T`). Hydrogens were regenerated by `pdb2gmx -ignh`.
- **Ion parameters.** Standard Åqvist-type AMBER monovalent and divalent parameters: $K^+$ $\sigma = 0.303797$ nm, $\varepsilon = 0.810369$ kJ mol$^{-1}$; $Cl^-$ $\sigma = 0.447766$ nm, $\varepsilon = 0.148913$ kJ mol$^{-1}$; $Mg^{2+}$ $\sigma = 0.276980$ nm, $\varepsilon = 0.0123428$ kJ mol$^{-1}$.
- **Ion counts.** Added by `genion` in three passes by replacement of water: (i) 196 $K^+$ for neutralisation, (ii) 165 $K^+$/165 $Cl^-$ for 0.15 M KCl, (iii), 5 mM $Mg^{2+}$

- **Integration.** Leap-frog integrator, time step $\Delta t = 2$ fs in all dynamics stages except NVT ($\Delta t = 1$ fs); bonds to hydrogen constrained with LINCS (`lincs_order` 4, `lincs_iter` 1).

- **Neighbour searching.** Verlet buffered lists, grid-based, `nstlist` = 200, buffer tolerance $5\times10^{-3}$ kJ mol$^{-1}$ ps$^{-1}$ per atom, periodic boundary conditions in $x$, $y$, $z$.

- **Electrostatics.** Particle-mesh Ewald, real-space cutoff $r_{\text{coulomb}} = 1.0$ nm, interpolation order 4, requested Fourier spacing 0.12 nm (actual grid $120 \times 120 \times 120$, 0.114 nm).

- **Van der Waals.** Cutoff $r_{\text{vdw}} = 1.0$ nm with potential-shift (Verlet) modifier and long-range dispersion corrections applied to both energy and pressure (`DispCorr = EnerPres`).

- **Temperature coupling.** Velocity-rescaling thermostat on two groups, `Solute` and `Water_and_ions`, with $\tau_T = 0.1$ ps for each and $T_{\text{ref}} = 310$ K for each. Initial velocities were drawn from a Maxwell–Boltzmann distribution at 310 K (or 333K depending the condition) with a random seed at the NVT stage only.

- **Pressure coupling.** Isotropic coupling to $p_{\text{ref}} = 1.0$ bar with $\tau_p = 2.0$ ps, isothermal compressibility $4.5 \times 10^{-5}$ bar$^{-1}$ and `refcoord_scaling = com`: Berendsen barostat during NPT equilibration, Parrinello–Rahman barostat during production.

- **Energy minimisation.** (i) Steepest descent during ion placement ($F_{\max} < 1000$ kJ mol$^{-1}$ nm$^{-1}$, step 0.01 nm, up to $5 \times 10^5$ steps); (ii) steepest descent of the ionised system (same tolerance and step, up to $2 \times 10^6$ steps, h-bond constraints); (iii) conjugate-gradient refinement ($F_{\max} < 100$ kJ mol$^{-1}$ nm$^{-1}$, step 0.01 nm, limited to 10 steps).

- **Simulation lengths.** NVT $5 \times 10^5$ steps = 500 ps; restrained-stage NPT $5 \times 10^5$ steps = 1 ns; extended NPT $5 \times 10^6$ steps = 10 ns; production $2.5 \times 10^8$ steps = 500 ns. Production output: coordinates to XTC every 25 000 steps (50 ps), energies and log every 10 000 steps (20 ps), no TRR output.

The available continuous production segments span 312.25–500 ns each. All primary comparisons use the common segment-local 0–300 ns interval. Table S6 records the systems, substitutions, simulation conditions, and production sampling included in the analysis.

#### S8.3. Trajectory processing

The simulations comprise four RNA sequences at two temperatures, giving eight sequence–temperature conditions and eleven continuous production segments (Table S6). Some simulations were interrupted and subsequently resumed from the endpoint reached before interruption. Equilibration was repeated before production sampling resumed. These restarted portions are continuations of the preceding simulations and share their history; they are not independent replicas. The reported averages combine the retained production sampling, while separate-segment results in Figs. S9 and S10 assess whether the differences persist before averaging.

Trajectories were analyzed with MDAnalysis 2.10.0 and Python 3.13.2 [15, 16], using aligned RNA coordinates. Each continuous production segment was assigned its own local time origin. Primary analyses use 0–300 ns of each segment. Frame pairs used for displacement calculations lie within a single production segment and never span an interruption or a repeated equilibration interval.

Backbone displacements were calculated from frames sampled every 0.25 ns. Pocket-distance calculations used every fifth saved frame, corresponding to the same nominal interval. The pocket-sampling stride was maintained across the trajectory, so the first sampled frame of a restarted segment can be offset from its local time origin by up to approximately 0.2 ns. Only frames within the specified analysis window were included. To assess sensitivity to the analysis window, we repeated the calculations over 100–300 and 200–300 ns and over the three disjoint 100 ns intervals. The latter intervals include their left endpoint and exclude their right endpoint, except that the final 300 ns endpoint is included.

For pocket distributions and means, each retained frame has equal weight. For mean whole-chain displacements, each valid within-segment frame pair has equal weight. Per-nucleotide RMS displacements are obtained by averaging squared residuals over the selected atoms and valid frame pairs before taking the square root. Thus, each segment contributes in proportion to its retained frame or frame-pair count, as appropriate. Separate-segment curves show how the pooled results relate to the individual segments.

TABLE S6. **MD systems, simulation conditions and available production sampling.** **a**: sequence identities and substitutions relative to Start–10, in full-construct numbering. WT differences are listed in the same direction, Start–10 to WT, for comparison. **b**: durations of the continuous production segments analyzed. A slash separates segments; it does not denote an uncertainty or establish independent replication. **c**: settings common to all systems. Interrupted simulations resumed from their previous endpoints after repeated equilibration, and their production portions are treated as continuation segments. Primary analyses use 0–300 ns in each segment's recorded local time.

**a. Systems and substitutions**

| | |
|---|---|
| WT | *Azoarcus* WT; 197 nucleotides. Differences from Start–10: U4C, G6U, G11C, G21A, U22C, U24C, C33G, U46C, A56G, U57C, U60C, G61C, A64C, A65C, C66G, A67G, A70G, C71U, G99C, A100C, A101U, A102G, U108A, G133C, U144C, G148C, U149G, A172G, C185U, G194C. |
| Start–10 | Starting designed sequence; 197 nucleotides. |
| Jointly favoured | `start10_lowDCA_lowSB_031`; 197 nucleotides. Substitutions: G11U, A26C, G29C, C33G, C71U, A100C, G133C, U144C, A145C. |
| DCA favoured | `start10_lowDCA_highSB_007`; 197 nucleotides. Substitutions: U4C, G11C, C12U, U22C, C33U, C59A, G68A, C71U, C185U. |

**b. Available production segments**

| System | $T$ (K) | Segments | Durations (ns) | Window/segment (ns) |
|---|---|---|---|---|
| WT | 310 | 2 | 371.50 / 401.05 | 0–300 |
| Start–10 | 310 | 2 | 500.00 / 390.00 | 0–300 |
| Jointly favoured | 310 | 1 | 377.70 | 0–300 |
| DCA favoured | 310 | 1 | 500.00 | 0–300 |
| WT | 333 | 1 | 500.00 | 0–300 |
| Start–10 | 333 | 1 | 421.20 | 0–300 |
| Jointly favoured | 333 | 2 | 312.25 / 324.35 | 0–300 |
| DCA favoured | 333 | 1 | 500.00 | 0–300 |

**c. Common simulation conditions**

| | |
|---|---|
| Software | GROMACS 2025.1 |
| Force-field distribution | `amber14sb_99bsc0-cOL3_OPC` |
| Solvent and nominal salts | OPC water; 0.15 M KCl; 5 mM $MgCl_2$ |
| Periodic box | Dodecahedral; minimum clearance 1.2 nm |
| Thermostat / barostat | V-rescale / Parrinello–Rahman |
| Constraints | LINCS on bonds involving hydrogen |
| Equilibration | 0.5 ns NVT; 1 ns NPT; 10 ns unrestrained |
| Restart handling | Resume from previous endpoint; repeat equilibration; analyze subsequent production as a continuation segment |

#### S8.4. Whole-chain and per-nucleotide displacements

To measure changes in backbone conformation, we selected the phosphorus atom P and the sugar–phosphate backbone atoms C5′, C3′, O3′, and O5′ from each nucleotide. Five atoms per nucleotide would give $197 \times 5 = 985$ atoms. The first nucleotide, at the 5′ end, has no P atom in the simulated construct and contributes the other four atoms, giving $N_{\mathrm{bb}} = 984$. Each selected atom contributes equally to the structural fit and displacement calculation, without mass weighting. Coordinates and displacements are expressed in angstroms (Å), with 10 Å = 1 nm. Base atoms are used separately for the pocket-distance analysis below.

For a pair of frames separated by $\tau = 10$ ns, both coordinate sets are centered and optimally superposed with a proper least-squares rotation over the complete backbone selection. Let $\delta\mathbf{r}_a(t;\tau)$ be the resulting residual between atom $a$ in

TABLE S7. **The ten pocket residues and their reference correspondence.** Residue numbering in the first column refers to the full 197-nucleotide construct. Identities in that column are shared by WT, Start–10, the jointly favoured variant, and the DCA favoured variant. The 9V7P correspondence uses PDB chain and residue numbering. The final column gives the minimum distance between any heavy atom of that residue and any heavy atom of terminal ΩG in 9V7P; ΩG itself is included with distance zero.

| Construct residue | 9V7P chain:residue | Minimum distance (Å) |
|---|---|---|
| A118 | B:A126 | 2.838 |
| G119 | B:G127 | 2.541 |
| G121 | B:G129 | 2.926 |
| A122 | B:A130 | 4.887 |
| A163 | B:A171 | 3.129 |
| U164 | B:U172 | 2.534 |
| U167 | B:U175 | 4.374 |
| C168 | B:C176 | 3.137 |
| G196 | B:G204 | 1.602 |
| G197 (ΩG) | B:G205 | 0.000 |

frame $t$ and its counterpart in frame $t-\tau$. The whole-chain coordinate RMSD is

$$C_\tau(t) = \sqrt{\frac{1}{N_{\mathrm{bb}}}\sum_{a=1}^{N_{\mathrm{bb}}} \|\delta\mathbf{r}_a(t;\tau)\|^2}. \tag{S19}$$

Both endpoints must lie within the same segment and analysis window. At the sampling interval used here, 10 ns corresponds to 40 sample intervals.

For nucleotide $i$, with $n_i$ selected backbone atoms, the RMS displacement in window $W$ is

$$R_i(\tau;W) = \sqrt{\frac{1}{N_W n_i}\sum_{t\in\mathcal{P}_W}\sum_{a\in i} \|\delta\mathbf{r}_a(t;\tau)\|^2}, \tag{S20}$$

where $\mathcal{P}_W$ is the set of eligible frame pairs across the included segments, indexed by their later endpoints, and $N_W = |\mathcal{P}_W|$. Every nucleotide uses the same global fit; nucleotides are not fitted independently. The profiles satisfy

$$\sum_{i=1}^{197} \frac{n_i}{N_{\mathrm{bb}}} R_i(\tau;W)^2 = \left\langle C_\tau(t)^2 \right\rangle_W . \tag{S21}$$

Figure S9 shows the profiles at both temperatures for the full and later windows. At 60 °C, the jointly favoured pooled profile is below both Start–10 and the DCA favoured variant at 165/197 positions over 0–300 ns, 164/197 over 100–300 ns, and 106/197 over 200–300 ns. Requiring both jointly favoured segments separately to be below both comparators gives 130/197 positions over the full window. For terminal G197, the full-window pooled displacements are 3.02, 9.99, 8.03, and 10.38 Å for WT, Start–10, jointly favoured, and DCA favoured, respectively. The two jointly favoured segments give 9.87 and 5.61 Å, showing appreciable segment variability.

#### S8.5. Pocket selection and residue mapping

The fixed pocket ∂ΩG contains terminal guanosine ΩG and every nucleotide represented in the simulations with at least one heavy atom within 5 Å of an ΩG heavy atom in PDB 9V7P [17]. The selection is made once from the experimental structure and held fixed across systems, temperatures, segments, and windows. The cutoff applies to heavy-atom distances, not to base-centroid distances. It selects ten nucleotides and therefore $\binom{10}{2} = 45$ unordered pairs (Table S7). Nearby residues, ligands, ions, or waters absent from the simulated RNA construct do not add positions to the selection.

For chain B of 9V7P, the PDB residue number equals the simulation position plus eight; simulated G197 corresponds to G205. All 197 mapped WT identities agree with 9V7P. Neither designed derivative introduces a substitution within the pocket, and all four simulated sequences have the same identity at each pocket position.

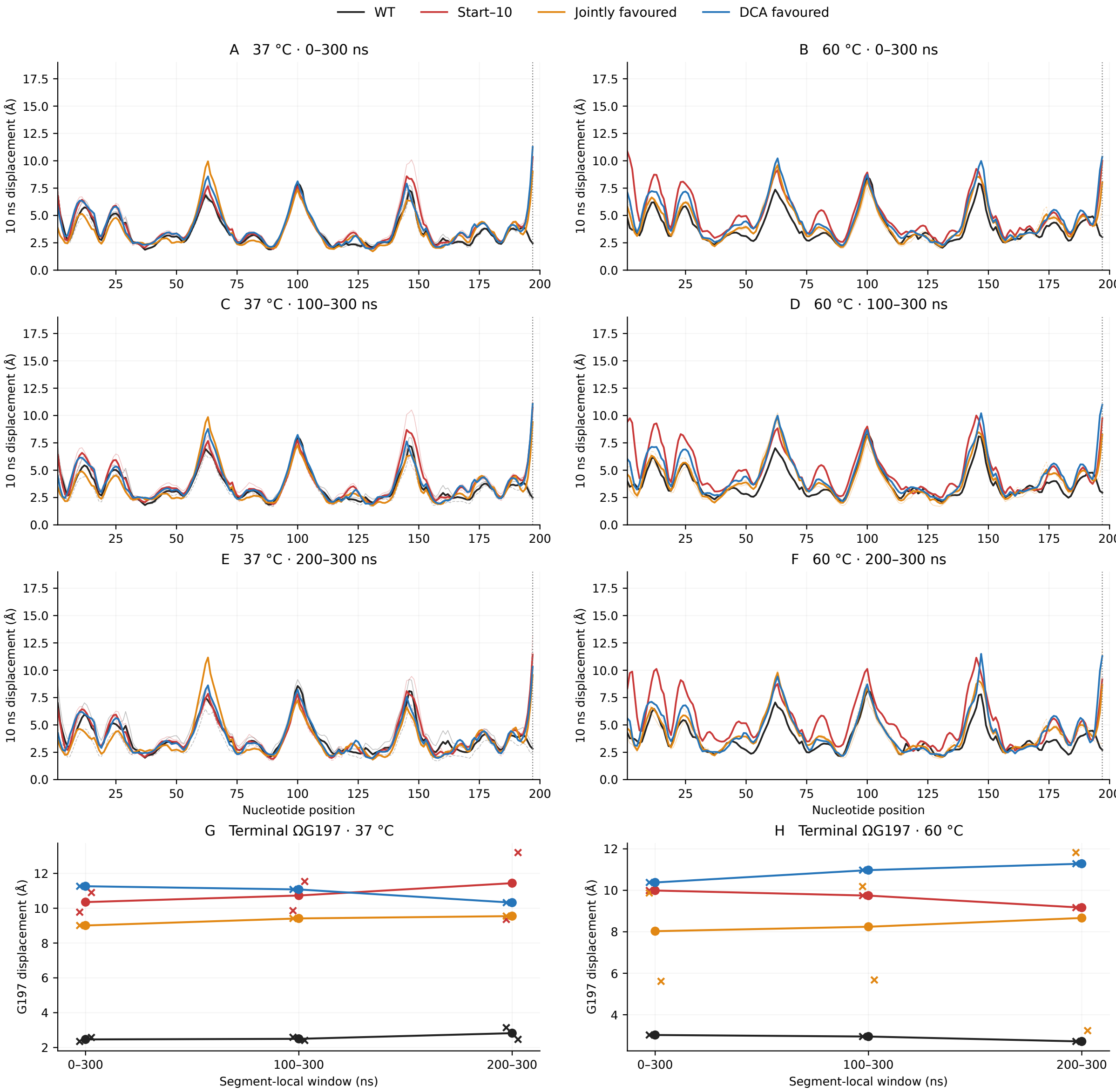


FIG. S9. **Sequence-wide displacements and analysis-window sensitivity. A–F**: per-nucleotide RMS displacements between globally fitted frames separated by 10 ns, at 37 and 60 °C, for segment-local windows 0–300, 100–300, and 200–300 ns. Thick curves pool squared residuals over valid within-segment frame pairs before taking the square root; thin curves show individual segments, with dashed lines for second segments. **G**,**H**: terminal ΩG197 displacements for the same windows; connected points are pooled values and crosses are segment-specific values. WT is black, Start–10 red, jointly favoured orange, and DCA favoured blue. All residues use the same global backbone fit. These quantities measure lagged displacement rather than RMSF.

#### S8.6. Pocket-distance metrics and structural insets

For nucleotide $i$, the base position is its unweighted heavy-atom centroid $\mathbf{c}_i(t)$. Hydrogen, sugar, and phosphate atoms are excluded. The standard-base atom sets are N9, C8, N7, C5, C6, N6, N1, C2, N3, C4 for A; N9, C8, N7, C5, C6, O6, N1, C2, N2, N3, C4 for G; N1, C2, O2, N3, C4, N4, C5, C6 for C; and N1, C2, O2, N3, C4, O4, C5, C6 for U. The inter-base distance is $d_{ij}(t) = \|\mathbf{c}_i(t) - \mathbf{c}_j(t)\|$, and $d_{ij}^{\mathrm{ref}}$ is the corresponding experimental distance. The experimental reference is 9V7P. These distances are invariant under a common rigid-body translation or rotation, so no superposition is required for this score.

With the main-text notation, the frame-level pocket distance RMSD is

$$D_{\partial\Omega\mathrm{G}}(t) = \sqrt{\frac{1}{45} \sum_{i,j\in\partial\Omega\mathrm{G}} \left(d_{ij}(t) - d_{ij}^{\mathrm{ref}}\right)^2}. \tag{S22}$$

Pocket-pair sums run over the 45 unique unordered pairs, counting each pair once. Its empirical cumulative distribution is

$$F(x) = \frac{100}{N} \sum_{t=1}^{N} \mathbf{1}\{D_{\partial\Omega\mathrm{G}}(t) \leq x\}, \tag{S23}$$

where $N$ is the retained frame count and $x$ is in Å.
For a pair of pocket residues, define the mean-squared distance deviation

$$\mathrm{MSD}_{ij} = \left\langle \left(d_{ij}(t) - d_{ij}^{\mathrm{ref}}\right)^2 \right\rangle. \tag{S24}$$

The main-figure maps display $\Delta\mathrm{MSD}_{ij} = \mathrm{MSD}_{ij,\mathrm{variant}} - \mathrm{MSD}_{ij,\mathrm{Start-10}}$, in Å$^2$, with each average evaluated at the same temperature and window. Negative values indicate smaller reference-distance deviations than Start–10. The maps do not report contact frequencies.
The per-residue distance RMSD used to colour the main-figure structural insets is

$$Q_i = \sqrt{\frac{1}{9} \sum_{\substack{j\in\partial\Omega\mathrm{G} \\ j\neq i}} \mathrm{MSD}_{ij}}, \qquad i \in \partial\Omega\mathrm{G}. \tag{S25}$$

Each residue has nine partners, with equal weight. The displayed inset value is

$$\Delta Q_i = Q_{i,\mathrm{variant}} - Q_{i,\mathrm{Start-10}}, \tag{S26}$$

in Å. The square root is taken separately for each sequence before subtraction. This is a distance-based residue summary, distinct from a Cartesian atomic RMSD. The insets display the variant models coloured with trajectory-derived $\Delta Q_i$ values; they are not selected trajectory snapshots. For consistency, these summaries obey

$$\frac{1}{10} \sum_{i\in\partial\Omega\mathrm{G}} Q_i^2 = \langle D_{\partial\Omega\mathrm{G}}^2 \rangle = \frac{1}{45} \sum_{i,j\in\partial\Omega\mathrm{G}} \mathrm{MSD}_{ij}. \tag{S27}$$

#### S8.7. Segment and time-window sensitivity

Figure S10 shows separate-segment and pooled pocket distributions, and segment-specific means for the five analysis windows, all relative to 9V7P. At 60 °C over 0–300 ns, pooled mean pocket RMSDs are 3.77 Å for WT, 11.20 Å for Start–10, 7.80 Å for jointly favoured, and 11.31 Å for DCA favoured. Both jointly favoured segments have smaller means than Start–10 and the DCA favoured variant in all five windows. Their full-window medians differ, at approximately 9.48 and 5.98 Å, so the direction is more consistent than the precise magnitude of the difference.

#### S8.8. Mean-geometry shifts and fluctuations

Reference-distance deviations combine differences in average geometry and temporal fluctuations. For each pair, the exact identity

$$\mathrm{MSD}_{ij} = \left(\langle d_{ij}\rangle - d_{ij}^{\mathrm{ref}}\right)^2 + \left\langle \left(d_{ij} - \langle d_{ij}\rangle\right)^2 \right\rangle \tag{S28}$$

separates these contributions. The first term is the squared deviation of the mean distance from the reference; the second is the within-segment distance variance. We evaluate both terms separately within each continuous segment over 0–300 ns and average over the 45 pairs, using population variance with divisor $N$. Their sum is exactly $\langle D_{\partial\Omega\mathrm{G}}^2\rangle$ for

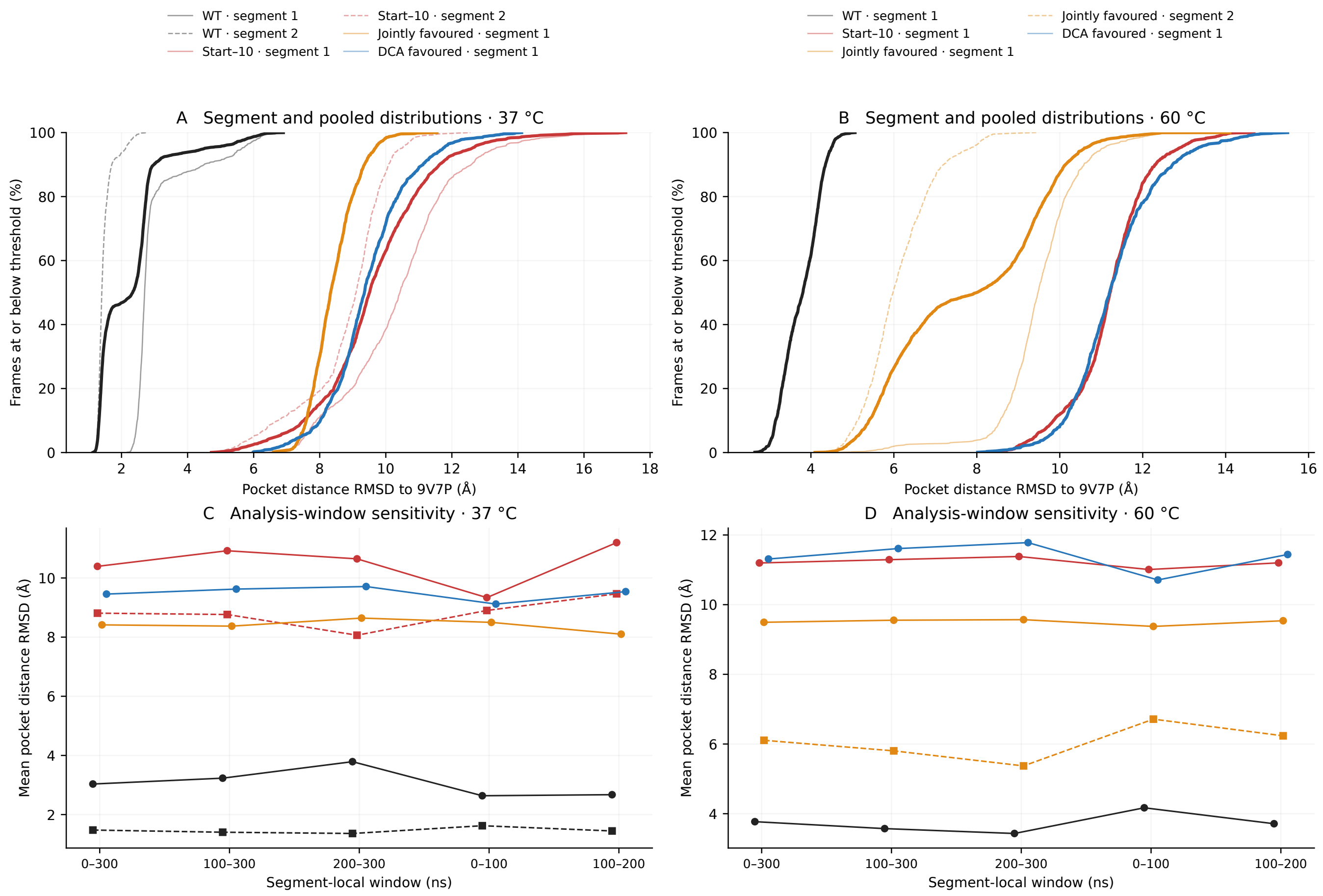


FIG. S10. **Pocket-distance sensitivity to continuation segments and analysis windows.** **A**,**B**: cumulative percentages of frames with pocket distance RMSD relative to 9V7P at or below the indicated threshold over 0–300 ns, at 37 and 60 °C. Thin curves show individual continuous production segments, with second segments dashed; thick curves pool retained frames. **C**,**D**: segment-specific mean scores over the 0–300, 100–300, 200–300, 0–100, and 100–200 ns windows. Circles and solid lines identify segment 1; squares and dashed lines identify segment 2. WT is black, Start–10 red, jointly favoured orange, and DCA favoured blue. Interrupted simulations resumed from their previous endpoints with equilibration repeated before production; the segments are continuations rather than independent replicas. All pocket scores use the same 45 unique pairs.

that segment. It does not decompose $\langle D_{\partial\Omega\mathrm{G}}\rangle$. Pooling segments before centering would include differences between segment means in the fluctuation term.

At 60 °C, squared mean-geometry deviations account for 92.8% of the WT error, 97.6% of the Start–10 error, 97.1% and 89.2% of the two jointly favoured segment errors, and 95.6% of the DCA favoured error. The corresponding pair-averaged distance variances are 1.04, 3.05, 2.69, 4.11, and 5.71 $\text{Å}^2$, respectively. In particular, the second jointly favoured segment has a lower reference error but larger distance variance than Start–10. The smaller pocket score therefore primarily reflects a more reference-like mean geometry, rather than uniformly reduced fluctuations.

#### S8.9. Displacements outside mutation neighbourhoods

To test whether the sequence-wide displacement difference extends beyond substituted positions, we exclude nucleotides within sequence distance $k$ of a substitution, with $k = 0$, 5, or 10. For jointly favoured versus Start–10, the mutation set $\mathcal{M}$ contains the nine jointly favoured substitutions. For jointly favoured versus DCA favoured, it is the union of the substitutions in either derivative relative to Start–10. This conservative rule also excludes positions carrying the same substitution in both derivatives. A construct position $i$ is retained when

$$i \in \mathcal{I}_k \quad \Longleftrightarrow \quad \min_{m \in \mathcal{M}} |i - m| > k, \qquad 1 \le i \le 197. \tag{S29}$$

These are sequence neighbourhoods, not spatial shells. The exclusion selects positions from the profiles already calculated with the full global backbone fit; the fit itself is not recomputed.
At 60 °C over 0–300 ns, excluding substituted positions alone leaves 167/188 retained sites with lower jointly favoured displacement than Start–10, and 163/182 with lower displacement than DCA favoured. Excluding ±5 nucleotides gives 104/123 and 78/91, respectively; excluding ±10 gives 65/79 and 39/46. These results support differences extending beyond the immediate mutation neighbourhoods.

---

[7] J. Abramson, J. Adler, J. Dunger, R. Evans, T. Green, A. Pritzel, O. Ronneberger, L. Willmore, A. J. Ballard, J. Bambrick, S. W. Bodenstein, D. A. Evans, C.-C. Hung, M. O'Neill, D. Reiman, K. Tunyasuvunakool, Z. Wu, A. Žemgulytė, E. Arvaniti, C. Beattie, O. Bertolli, A. Bridgland, A. Cherepanov, M. Congreve, A. I. Cowen-Rivers, A. Cowie, M. Figurnov, F. B. Fuchs, H. Gladman, R. Jain, Y. A. Khan, C. M. R. Low, K. Perlin, A. Potapenko, P. Savy, S. Singh, A. Stecula, A. Thillaisundaram, C. Tong, S. Yakneen, E. D. Zhong, M. Zielinski, A. Žídek, V. Bapst, P. Kohli, M. Jaderberg, D. Hassabis, and J. M. Jumper, Accurate structure prediction of biomolecular interactions with AlphaFold 3, Nature **630**, 493 (2024).
